# Geological degassing enhances microbial metabolism in the continental subsurface

**DOI:** 10.1101/2020.03.07.980714

**Authors:** Till L.V. Bornemann, Panagiotis S. Adam, Victoria Turzynski, Ulrich Schreiber, Perla Abigail Figueroa-Gonzalez, Janina Rahlff, Daniel Köster, Torsten C. Schmidt, Ralf Schunk, Bernhard Krauthausen, Alexander J. Probst

**Affiliations:** Environmental Microbiology and Biotechnology, Faculty of Chemistry, University Duisburg-Essen, Germany; Department of Geology, University Duisburg-Essen, Germany; Instrumental Analytical Chemistry and Centre for Water and Environmental Research (ZWU), University of Duisburg-Essen, Germany; Geyser-Center, Andernach, Germany; Institute of Applied Geosciences, Karlsruhe Institute of Technology, Germany

## Abstract

Mantle degassing provides a substantial amount of reduced and oxidized gases shaping microbial metabolism at volcanic sites across the globe, yet little is known about its impact on microbial life under non-thermal conditions. Here, we characterized deep subsurface fluids from a cold-water geyser driven by mantle degassing using genome-resolved metagenomics to investigate how the gases impact the metabolism and activity of indigenous microbes compared to non-impacted sites. While species-specific analyses of Altiarchaeota suggest site-specific adaptations and a particular biogeographic pattern, chemolithoautotrophic core features of the communities appeared to be conserved across 17 groundwater ecosystems between 5 and 3200 m depth. We identified a significant negative correlation between ecosystem depth and bacterial replication, except for samples impacted by high amounts of subsurface gases, which exhibited near-surface activity. Our results suggest that geological degassing leads to higher nutrient flows and microbial activity in the deep subsurface than previously estimated.

## Introduction

The continental subsurface is a huge reservoir for life, hosting about 60% of all microorganisms on Earth^1,2^. Carbon, nitrogen and sulfur turnover by these microorganisms have a vast contribution to all biogeochemical cycles on the planet^3^. In addition to the great number of microorganisms, subsurface ecosystems can accommodate a large diversity of different bacteria and archaea^4,5,6^, with even single ecosystems containing representatives of almost all known bacterial phyla^4^. Subsurface ecosystems are categorized as either detrital or productive, depending on whether buried organic carbon or inorganic carbon are the main carbon sources of the community^7^. Since no light is available as an energy source in the deep biosphere, alternative electron donors to water like hydrogen or sulfide are used to fuel mostly anaerobic carbon fixation pathways such as the Wood-Ljungdahl pathway^7^. Subsurface lithoautotrophic microbial communities^8^ have been reported for many terrestrial ecosystems including the Fennoscandian shield^9^, the Columbia River basalt^8^, the Witwatersrand Basin^10^, and subsurface fluids discharged by Crystal Geyser^11^. While these subsurface ecosystems are usually dominated by bacteria, one exception are archaea belonging to the Alti-1 clade of the Altiarchaeota^12,5,13^. Alti-1 form biofilms using their characteristic nano-grappling hooks (*hami*)^14,15^. The other clade, Alti-2, are more widespread and diverse but found at lower abundances in their ecosystems^14^. Altiarchaeota are able to live autotrophically using the most energy efficient carbon fixation pathway^16^, which was the most dominant carbon fixation pathway prior to the evolution of photosynthesis^17,18^.

Chemolithoautotrophic life in subsurface ecosystems necessitates the presence of adequate electron donors like hydrogen, hydrogen sulfide or methane. One source of such gases can be Earth’s mantle, which also releases huge amounts of oxidized carbon, mainly in form of carbon dioxide (CO_2_), into the crust and the atmosphere^19^. This process, also termed mantle degassing, is the transition of volatiles from the mantle (supercritical) to the subcritical zone of the upper crust fueled by lower pressure of volatiles near the surface compared to the mantle^20^. Modern Earth has few areas with active mantle degassing, which are usually restricted to terrestrial volcanoes or hydrothermal vents in oceans^21,22,23,24^. At hydrothermal vents, chemolithoautotrophs initiate the microbial trophic network and proliferate at high rates leading to high microbial cell numbers^1^. While volcanic sites and hydrothermal vent fields have been studied fairly thoroughly regarding both their microbiome and their microbial activity^25,26,27,28,29^, little is known about deep subsurface ecosystems with low temperatures but heavy impact from gases released from the mantle.

Previous studies have analyzed the influence of mantle degassing via volcanic mofettes on near surface biomes, particularly soil microbial communities^30,31,32,33^. Mehlborn et al.^31^ showed that gasses from the mantle can alter the availability of different heavy metals including metalloid arsenic and predicted impacts on microbial communities. In 2015 and 2016, Beulig and co-workers reported an increase in dark carbon fixation and found evidence that the carbon dioxide from the degassing is indeed incorporated into biomass^32,33^. Along with fermentation processes, the pathways for the turnover of organic carbon were similar in both systems, while the microbial diversity of soils impacted by gases from the mantle was lower compared to controls. Carbon and sulfate respiration were enriched during degassing, while aerobic respiration declined^33^, and acetogenesis was suggested to play a major role in these systems^32^. However, these studies were limited to the upper 50-cm of Earth’s critical zone and the influence of mantle degassing on mesophilic microbial communities in the deep subsurface including their metabolic capacity and activity has not been investigated so far.

The cold-water Geyser Andernach is located in the Rhine Valley near Koblenz in western Germany and is driven by gases discharged from the mantle^19^. Since 2001 the geyser has had an intact tubing, thus tapping into a unique ecosystem. Once released by a mechanical shutter, the gasses from the mantle (mainly CO_2_) permeating the groundwater cause the eruption of cold subsurface fluids sourced from an uniform aquifer system. Thus, Geyser Andernach is an ideal ecosystem to investigate how mantle degassing shapes mesophilic microbial life in the subsurface. Here, we used a combination of long-term geochemical characterization coupled to genome-resolved metagenomics to investigate the geyser’s microbial community. To analyze how mantle degassing impacts mesophilic microbial communities, we determined a geographical pattern of the most abundant organism across multiple ecosystems on different continents and set the bacterial replication indices and microbial metabolism abundances in Geyser Andernach into relation to 16 other deep continental subsurface ecosystems.

## Results

### Geyser Andernach provides access to a stable ecosystem impacted by mantle degassing

Geyser Andernach was drilled to a depth of 351 m in 1903 tapping into a shale-hosted aquifer with quartz veins. Its eruptions are driven by mantle degassing and can be controlled via mechanical shutters. Geochemical measurements averaged over 14 years have demonstrated that the subsurface fluids provide a constant environment (Table S1). The gaseous and ionic composition of the geyser showed the predominance of CO_2_ in the system and previously reported traces of hydrogen and hydrogen sulfide^19^. Prominent electron donors and acceptors were determined to be hydrogen and ferric iron as well as sulfate, respectively. To investigate the community in subsurface fluids impacted by mantle degassing, we sampled two eruptions of Geyser Andernach, and collected microorganisms onto three individual 0.1-µm filters. Metagenomic sequencing of the community resulted in ∼7 billion bp per sample (5% SD) assembled into 921,520 scaffolds on average (20% SD, for further statistics please see Table S2). Approximately 75% of the reads (2.6% SD) mapped back the assembly providing evidence that the reconstructed metagenome is representative for the ecosystem. The community composition based on rpS3 sequences assembled from the metagenome displayed a fairly restricted diversity consisting of 52 organisms, which spanned twelve phyla (Fig. 1). The core community was composed of 15 organisms detected across all three metagenomes (Fig. 1), and they accounted for 42.8% (1.3% SD) of the total relative abundance of the detected community. For 20 of these 52 microorganisms, we reconstructed high quality genomes with at least 70% estimated completeness (and less than 10% estimated contamination, details in Table S3). Interestingly, the most abundant species recruited 42.8 (1.3% SD) of the metagenomic reads and belonged to the phylum Altiarchaeota^5^ (in the following denoted as Altiarchaeum GA) and specifically grouped within the Alti-1^14^ clade. The second most abundant organism was classified as Caldiserica, which were originally known to inhabit hot springs^34^ but were recently also detected in subsurface ecosystems populated by mesophiles^11,5^.

**Figure 1.**
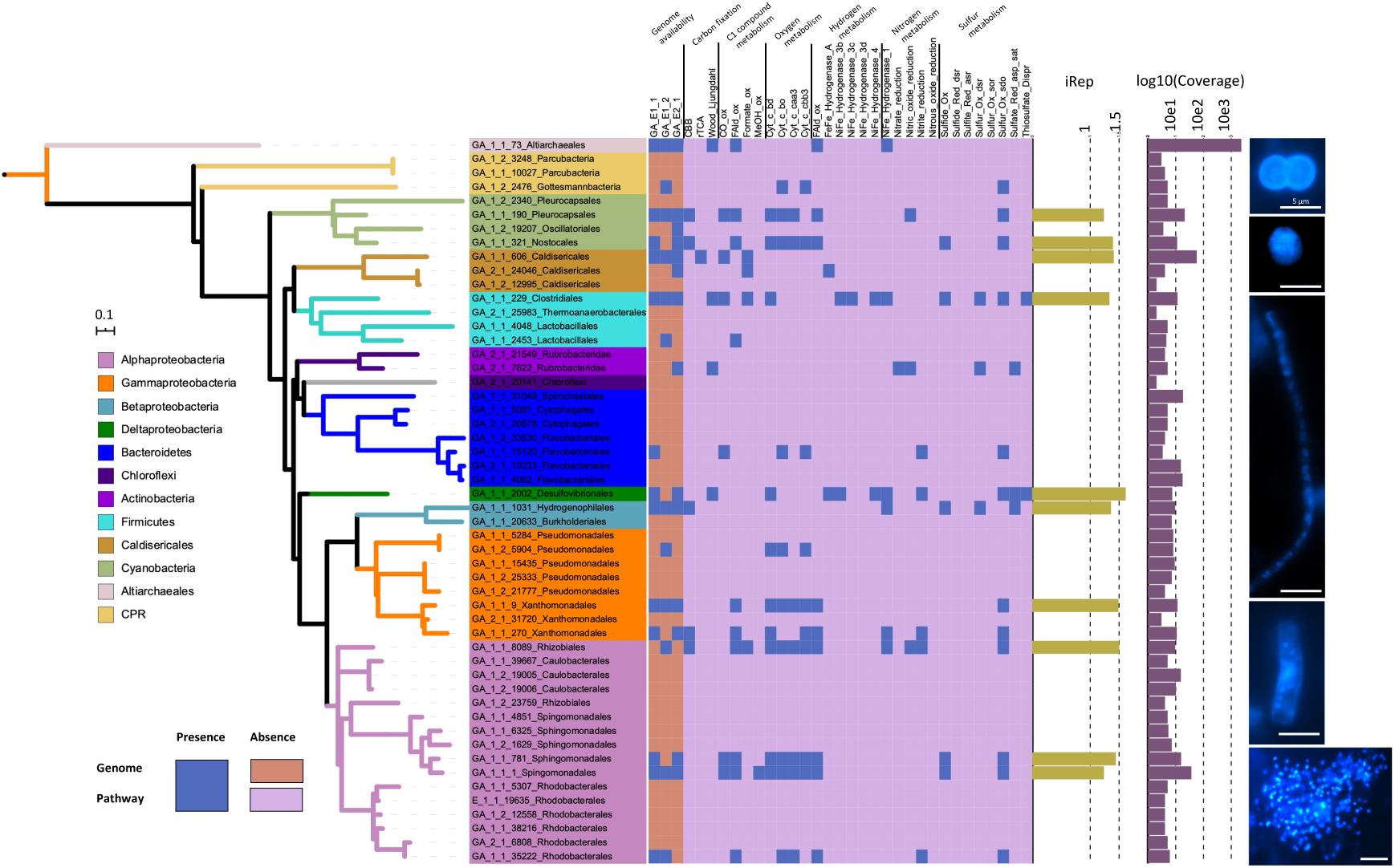
Metagenomic and microscopic characterization of the community in subsurface fluids discharged by Geyser Andernach. **A:** rpS3-based phylogenetic diversity of the organisms in the Geyser Andernach. Centroid rpS3 sequences (after clustering at 99% similarity using cdhit) were used for the calculation of the phylogenetic tree using IQTree. Colors of the different branches signify different phyla. The presence of recovered draft genomes in the three metagenomes (column “genome availability”) as well as the presence of marker genes for specific chemolithoautotrophic pathways is provided as blue boxes, with brown / pink boxes representing their absence. Violet bars show the mean relative log10-scaled abundance and olive bars show the average iRep value if multiple genomes could be recovered, the single iRep value if only one genome could be recovered. **B:** Morphologies of microorganisms as determined via DAPI staining and fluorescence microscopy (scale bars = 5 µm).

We estimated the activity of the bacteria in the community using *in situ* replication indices as an approximation. These indices ranged between 1.4-1.5, indicating that 40-50% of the analyzed bacterial population were undergoing genome replication at the time of sampling. Microscopic cell counts of organisms from the subsurface fluids ranged from 2.7 × 10^6^ to 4.2 × 10^6^ (average 3.5 × 10^6^) cells ml^-1^ (Fig. S1) and displayed various morphologies ranging from cocci and rods to filamentous-shaped microorganisms (Fig. 1). Importantly, we also observed clusters of small cocci, which are similar to previously reported biofilm structures of Altiarchaeota^12^ and whose presence is in agreement with the metagenomic results. We estimated the total amount of erupting carbon (CO_2_ and HCO_3_^-^) to be 6.27 t per year, while the microbial cells account for approximately 0.72 kg of carbon, suggesting that about 0.01% of carbon degassing from the mantle is fixed in this ecosystem.

### Biogeography and functional adaptations of deep subsurface Altiarchaota

Altiarchaeota can be divided into two clusters, Alti-1 and Alti-2, with the latter having a broader metabolic variability than Alti-1^14^. However, organisms of the Alti-1 cluster are those that can dominate entire ecosystems, as shown for multiple ecosystems across the globe^5,12,13,35^. We investigated the metabolic capacities of the newly recovered Altiarchaeum GA genome in comparison with other Altiarchaeota (Fig. 2A/B, details in Table S4), some of which we newly reconstructed from public datasets. We identified that all of them share a central NAD(P)H-based Wood-Ljungdahl pathway for carbon fixation and carbon monoxide utilization. The main difference of Altiarchaeum GA to most other deep subsurface Altiarchaeota was the presence of genes for a NiFe hydrogenase (Fig. 2B), which seems to be a specific adaptation to hydrogen containing gases from the mantle. This class of hydrogenases has so far only been detected in organisms of the Alti-2 group^14^ and in an Alti-1 genome from the Alpena Fountain^35^. Additionally, Altiarchaeota employ a strictly anaerobic metabolism with only few members possessing an oxygen stress response system^12^, likely not enabling them to be aerobically dispersed across larger distances.

**Figure 2.**
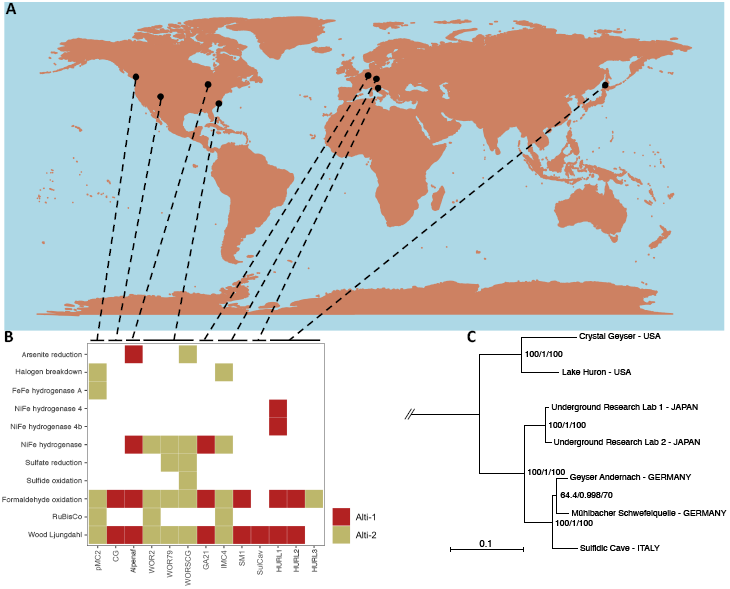
Geographical distribution and chemolithoautotrophic potential of Altiarchaeota. **A:** Global map with locations from which Altiarchaeota genomes were recovered. **B:** Metabolic potential of Altiarchaeota genomes. Alti-1 genomes are highlighted in red, Alti-2 genomes in green. **C:** Phylogeny of Alti-1 genotypes based on 30 universal ribosomal proteins (5136 aa positions, IQTree JTTDCMut+F+G4) and using the Alti-2 genome IMC4 as the outgroup. Branch supports correspond to ultrafast bootstraps^69^ (1000 replicates), the SH-aLRT test^70^ (1000 replicates), and the approximate Bayes test^86^ respectively (tree with outgroup in Fig. S3). Details on Altiarchaeales genomes in Table S4.

In contrast to these site-specific adaptations, the average nucleotide (ANI) and amino acid (AAI) identity of all so-far recovered Alti-1 indicated that they belong to the same genus (Fig. S2), although 16S ribosomal RNA gene similarity suggested the same species. When correlating the genomic differences based on ANI to the geographical distance between sampling sites of the Alti-1 genomes, a highly significant negative correlation (*Pearson*, cor = -0.77, p = 9 x10^−4^) could be observed, indicating that a greater distance led to greater dissimilarity (Fig. S3). We challenged this observation by using robust phylogenetic analyses based on a supermatrix of 30 ribosomal proteins and found that Altiarchaeota cluster based on geographical sampling site going all the way to continent scale (Fig. 2C). This indicates an extreme degree of biogeographic provincialism across Earth. The small genetic divergence of Alti-1 organisms in their core genome combined with their previously determined constant cell division^12^ implies a very slow evolutionary rate of these organisms in deep subsurface environments.

### Replication indices across multiple deep continental subsurface ecosystems

To investigate if mantle degassing has an impact on microbial activity in the continental subsurface, we used *in situ* replication indices (iRep) of bacterial genomes. We first investigated if iRep can be used as a measure of activity by comparing groundwater fluids to sediments, whereas sediments are known to be more active^36^. Indeed, iRep suggested a significantly higher activity in sediments than groundwater. Replication measures from Geyser Andernach were then compared with those from other public datasets from deep subsurface environments of varying depth (overview of samples and ecosystems is provided in Table S5). The sampling depth varied from 0 m below ground (cave systems) down to 3140 m depth. We reconstructed genomes of previously unbinned metagenomes resulting in 572 newly assembled and classified bacteria (Table S3) representing 415 different organisms after dereplication. Combined with genomes and iRep results from previous studies^5,4,13^, we leveraged *in situ* replication measures of 895 bacteria (Table S6) spanning the vast majority of all known bacterial phyla (see Supplementary File S1). Interestingly, the average iRep value of bacteria of the individual ecosystems correlated negatively and highly significantly with sample depth when using all iRep values individually (p-value < 10^−8^) and also when using the median per sampled ecosystem (p-value < 0.0007, Fig. 3, Table S7). In other words, the deeper the origin of the retrieved sample, the lower the genome replication measure.

**Figure 3.**
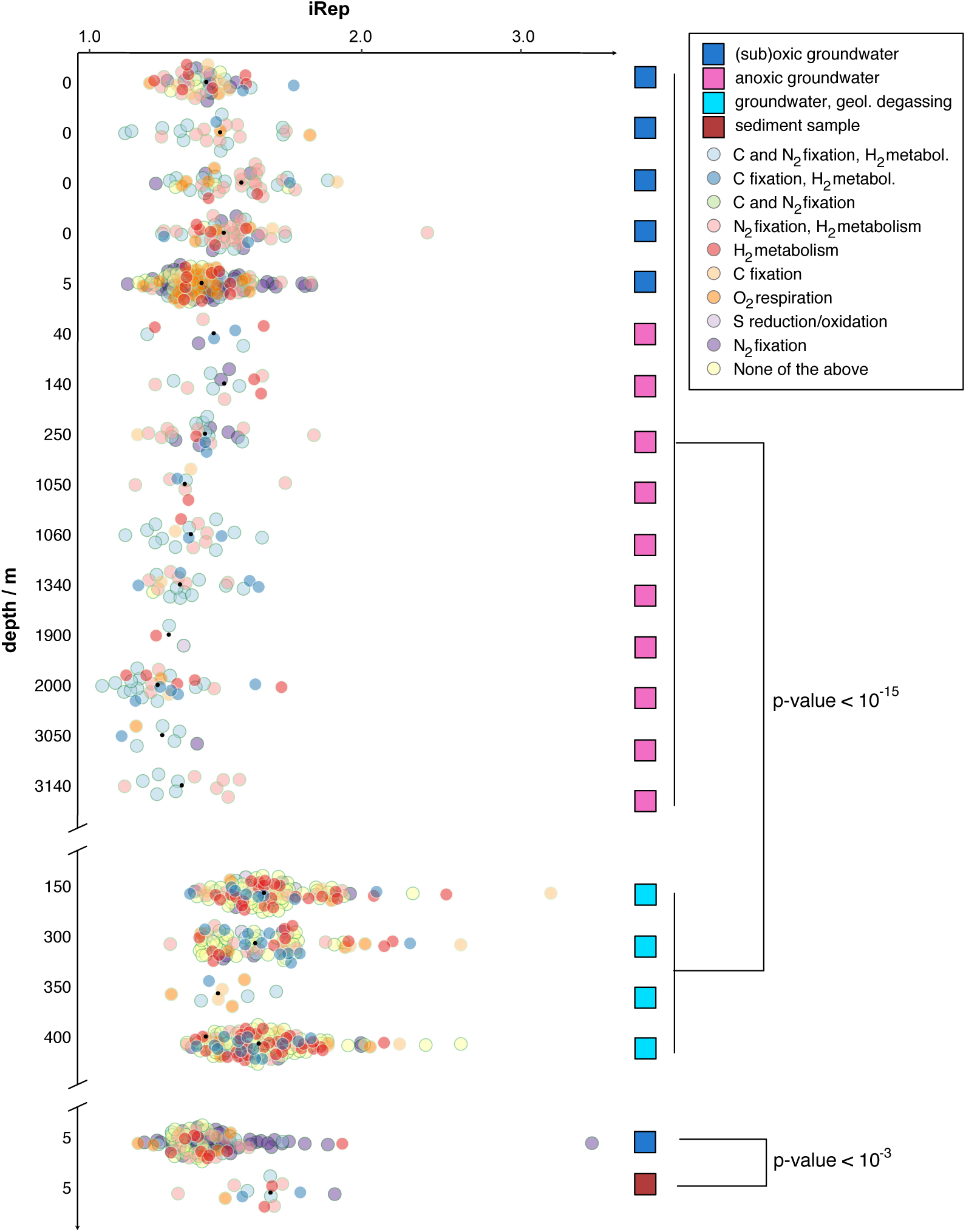
In situ bacterial replication rates across subsurface ecosystems ordered by ecosystem depth. The figure depicts a beeswarm plot of iRep values of genomes (x-axis) across ecosystems (y-axis) with genomes colored according to their predicted metabolic potential and the black dot representing the median iRep value (individual iRep values in Table S6). Samples impacted by geological degassing and a sediment sample along with the respective aquifer sample are plotted separately. The top y-axis shows the sampling depth of the different ecosystems (Table S7). In total, 895 genomes were used for this analysis with >=70% completeness and <=10% contamination based on 51 bacterial and 38 archaeal single copy genes. The order of samples is given in Table S7.

In particular, organisms with the capacity of carbon fixation (−0.47), of sulfur oxidation (−0.46) or of metabolizing hydrogen (−0.45) contributed to this observation (correlations are summarized in Table S8). Samples impacted by high CO_2_ concentrations, either solely from mantle degassing (this study) or from both mantle degassing and thermal activity^5^, were outliers in this correlation analysis. In fact, iRep measures of bacteria in these samples were significantly higher than iRep measures of other subsurface samples (p-value < 10^−15^) and nearly reached values of samples that are close to Earth’s surface (Fig. 3). When excluding these samples from the correlation analysis with depth, the respective correlation coefficient increased from -0.20 to -0.28 (p-value < 10^−8^). We also tested how the availability of oxygen influences genome replication measures of bacteria in the continental subsurface. iRep values were on average 0.09 higher for bacteria in oxygenic samples (p-value < 10^−8^) meaning that about 9% more of the bacteria were undergoing genome replication.

### Conserved chemolithoautotrophic metabolism of subsurface microorganisms

Since Altiarchaeota appeared to have a very specific adaptation to their respective ecosystems, we 234 investigated if the general metabolism for carbon, nitrogen and sulfur turnover of entire communities is adapted to high-CO_2_ subsurface environments. We searched for key enzymes for metabolic pathways across our entire metagenomic assemblies (Table S2) and used the abundance of scaffolds that carried a key enzyme as relative abundance measure of the respective metabolism (Fig. 4). Interestingly, the core metabolism remained relatively stable across all tested ecosystems. We performed both students t-tests and Kruskal-Wallis-tests along with equivalence testing to determine whether there was a significant difference between high-CO_2_ and non-high-CO_2_ metabolisms and could only detect a significant difference in the nitrite reduction metabolism (Kruskal-Wallis group comparison p-value = 6 × 10^−4^, details on tests in Table S9). Consequently, there seems to be very little difference in the metabolic potential between regular subsurface microbial communities and those at sites impacted by mantle degassing, although the indigenous organisms at these sites appear to have higher activities based on replication indices.

**Figure 4.**
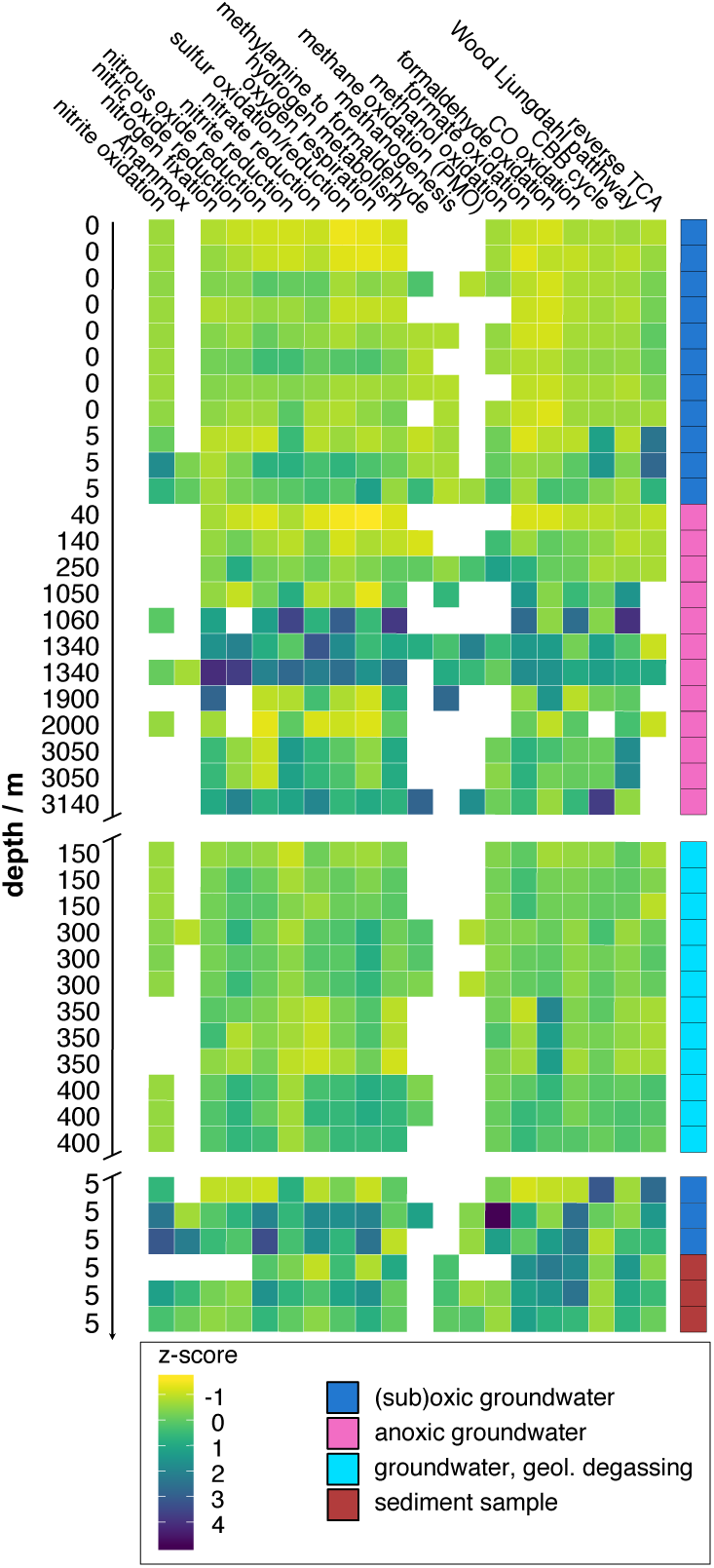
Chemolithoautotrophic metabolic potential across ecosystems. The heatmap shows the read-normalized abundance of chemolithoautotrophic pathways, Z-score scaled for the respective metabolisms. If multiple biological replicates of samples were available, up to three were depicted. Sample order is according to Fig. 3 and Table S7.

## Discussion

Modelling of current cell counts estimate the amount of prokaryotic microorganisms in the continental subsurface to 2 to 6 × 10^29 1^ which amounts to 60% of the prokaryotic life on our planet^2^. Interestingly, the diversity of microorganisms declines with sampling depth in the continental subsurface^1^. Our metagenome assemblies showed the same trend in diversity change (based on the rpS3 marker gene, cor=-0.40, p-value = 0.021, Fig. S4). This indicates that they are representative of general subsurface microbial communities and were consequently used to establish a genome database to calculate genome replication indices across various subsurface ecosystems. Estimation of genome replication indices is based on the measurement of the amount of DNA attributable to the origin versus the terminus of replication^37^, which indicates the presence of replication forks. In nutrient rich ecosystems, however, polyploidy can be a strategy to store phosphate^38^ or to increase the cell size^39^, causing the replication measures to no longer unambiguously indicate active replication. In contrast, the activity to acquire organic carbon, cellular growth and genome replication should be in agreement in oligotrophic systems like the deep terrestrial subsurface. It has been well established that sediments act like seed banks for aquifer communities of bacteria^40,41^, and the activity of microbes in sediments has been estimated to be higher than that of planktonic microorganisms^36^. Our data provide evidence that sessile bacteria have significantly higher genome replication measures than bacteria within the very same aquifer^4^, while also containing a much more homogeneous community (Fig. S4). This finding validates that replication indices can be used as proxies of activity in these subsurface ecosystems. Using these measures across multiple samples from different depths established that the activity of bacteria declines with sampling depth, suggesting a deep biosphere with little activity.

Analyses of predicted key enzymes across 41 assembled metagenomes implies that there is little variation in the deep biosphere’s metabolic potential. This metabolic conservation is, however, independent of the composition and relative abundance of detected taxa enabling the conclusion of metabolic redundancy across a large diversity^4^, which is probably influenced by the geochemistry of each system^42^. Interestingly, Alti-1 Altiarchaeota also showed a conservation in their core genome depicting a strict biogeographic distribution with limited recombination. The relationship between genomic similarity and geographic distance in general is a well-known occurrence in nature and has led to the development of concepts like the isolation-by-distance theory^43,44,45,46^. Since members of the Alti-2 group of Altiarchaeota^14^ show no biogeographic isolation, isolation-by-distance seemed unlikely at least for some subsurface microorganisms. Altiarchaeota of the Alti-1 group are sensitive to oxygen but possess *ham*i (hook-like cell surface appendages) to anchor entire biofilms on various surfaces^15^ making plate tectonics mediated-dispersal and isolation the most plausible mechanism for the observed pattern (for more discussion on the dispersal of Altiarchaeota, see the Supplementary Discussion). Nevertheless, the core genome analysis of Alti-1 Altiarchaeota showed very specific adaptations to each ecosystem suggesting other major evolutionary processes (like horizontal gene transfer) of these archaea in the subsurface.

At Geyser Andernach, Altiarchaeota of the Alti-1 clade reach high cell densities in the CO_2_ subsurface ecosystem and represent the main primary producers similar to the other high CO_2_ aquifer system, the Crystal Geyser, which additionally harbors a tremendous amount of bacterial diversity but also taps into three different aquifer ecosystems^11,5^. Interestingly, replication indices of bacteria from high CO_2_ aquifers did not follow the above-mentioned decline of activity with increasing depth. Specifically, we had two samplings sites, the cold high-CO_2_ geysers Geyser Andarnach and Crystal Geyser, which enabled access to four different ecosystems of different depth and substantially different community composition^5^. In these ecosystems impacted by mantle degassing, the bacterial replication measures reached values similar to near surface communities. The most obvious explanation relates to the unique geology caused by the mantle and thermal degassing. In sediment-hosted aquifers, groundwater flows through pore space, which generally decreases with depth as the sediment becomes more compact and the pores mineralize. Consequently, the groundwater flow rate decreases with depth in sediment-hosted aquifers. Since the availability of nutrients is directly proportional to the flow of the groundwater, deeper aquifers should have less replication activity. In contrast, fracture-controlled aquifers, which are characterized by solid rock formation-embedded channels, allow for flows up to multiple magnitudes greater than comparable sediment-hosted aquifers. We conclude the increased microbial activity in ecosystems impacted by mantle degassing stems from a combination of the availability of reduced mantle gasses like H_2_ and H_2_S as microbial electron donors and the increase in nutrient flow in these aquifers. Transferring these results to Early Earth, where mantle degassing heavily shaped the surface of Earth, we suggest a scenario with high microbial activity at our planet’s surface, which might have had large implications on the proliferation and evolution of life in its early stages.

## Material and Methods

### Geological setting

The cold-water geyser of Andernach is located 2 km downstream of Andernach (Rhine kilometer 615) on a 21-hectare peninsula called Namedyer Werth in the Middle Rhine valley. Driven by magmatic carbon dioxide, the geyser erupts regularly and intermittently approx. every 2 hours, when the groundwater filling the well is saturated with CO_2_ and a reinforced chain reaction (domino effect) concludes in a gas/water-eruption up to >60 meters in height^47^, lasting for 15-20 minutes. The well (drilling Ø 750/312/216 mm; casing/screens Ø 150 mm) was drilled in 2001 and is the third borehole (after 1903 and 1955) on this peninsula. The drilling taps 14 m of Quaternary fluvial deposits and continues then until its total depth of 351.5 m in lower Devonian formation called ‘Hunsrück Schiefer s.l.’ (shale)^48^.

The small peninsula is part of the Pleistocene terrace which is covered by a thin sandy layer of fluvial Holocene deposits. Only at the NE margin of the peninsula the terrace is bare of deposits. The thickness of the Quaternary layer varies from 14 m (drilling 2001) to 20.75 m (drilling 1903)^49^ and 24.2 m (drilling 1955) in the vicinity of the cold-water geyser. Beneath the Quaternary deposits follow lower Devonian rock formations of low metamorphic shale, such as clayish shale and intercalated minor layers of quarzitic sandstones; the thickness of these series is up to 5000 m.

The peninsula is located in the Middle Rhine Valley, which is a part of the European Cenozoic Rift System^50^. This rift system runs between the cities Bingen and Bonn in SE-NW-direction and crosses the Variscian complex of the Rhenish massif. Located at the SE edge of the lower Middle Rhine Valley, Geyser Andernach is situated on the intersection of two major fault structures: about 1 km to the NW the Variscian Siegen thrust fault running SW-NE crosses the Rhine Valley and can be traced for over 100 km from the Eifel area to the Westerwald. This fault shows a vertical displacement of several thousand meters, which occurred during the Variscian orogenesis, thus bringing rocks of the middle Siegenian stage in lateral contact with lower Emsian stage^51^. About 2 km to the SE the lower Middle Rhine valley is morphologically separated from the adjacent intraplate Tertiary Neuwied basin by an approx. 100 m vertical displacement caused by the SW-NE trending Andernach fault.

The Andernach fault and the Siegen thrust fault were in post-Variscian time intersected and 200-300 m displaced by a SE-NW trending dextral strike-slip fault^52,53^. The fault is supposed in the river Rhine bed and covered by Quaternary deposits. The horizontal movement was probably combined with shear strain and cataclastic rocks in the vicinity of the fault. This fault is the cause for pathways of mantle gases to reach the subsurface aquifers and ultimately the atmosphere.

Starting in Tertiary, a mantle plume under the Eifel area caused an uplift of the Rhenish massif during the last two million years and is the driving force for the volcanic activity in the Quaternary Eifel area since 700 k years^54^.

The mantle plume is the basic requirement for the rise of magma under and into the crust, whereby magmatic gases are released.

### Sampling and geochemical measurements

The mesophilic and CO_2_-driven Geyser Andernach (50.448588°N, 7.375355°E) in western Germany was sampled on 21 February in 2018 by collection of erupting water in sterile, DNA-free containers and subsequent filtration onto 0.1 µm pore size filters of 142 mm diameter (Merck Millipore, JVWP14225) and storage on dry ice / -80 °C until DNA extraction. Water samples were collected during eruption of the geyser and analyzed biochemically as well as microscopically (see Suppl. Material for details). In total, two sequential eruptions were sampled, resulting in two filter samples for the first eruption and one filter for the second eruption.

### Metagenomic sequencing and processing

DNA was extracted from three individual 0.1 µm bulk water filtration filter membranes using the DNeasy PowerSoil DNA extraction Kit (Qiagen, JVWP14225) according to the manufacturer’s instructions and was sequenced as part the Census of Deep Life phase 13 sequencing grant using Illumina NextSeq (paired end, 150 bps each). Quality control of raw reads was performed using BBduk (Bushnell, https://jgi.doe.gov/data-and-tools/bbtools/bb-tools-user-guide/) and Sickle^55^. Reads were assembled into contigs and scaffolded using metaSPAdes 3.11^56^. For the sample IMS-BF, a sub-assembly of reads not mapping to the available Altiarchaeales SM1 genome (GCA_000821205.1) was performed to improve assembly quality and this sub-assembly was used for the binning of additional genomes. Open reading frames were predicted for scaffolds larger than 1kbp using Prodigal^57^ in meta mode and annotated using DIAMOND blast^58^ against UniRef100 (state Dec. 2017)^59^, which contained the NCBI taxonomic information of the respective protein sequences. Taxonomy of each scaffold was predicted by considering the taxonomic rank of each protein on the scaffold on each taxonomic level and choosing the lowest taxonomic rank when more than 50% of the protein taxonomies agree. Reads were mapped to scaffolds using Bowtie2^60^ and the average scaffold coverage was estimated along with scaffolds’ length and GC content.

### Binning of GA samples

Abawaca^37^, MaxBin2^61^, tetranucleotide-based Emergent Self-Organizing Maps (ESOM^62^), and CONCOCT^63^ were used to identify metagenome assembled genomes and DAS Tool with standard parameters was used to aggregate the results^64^. (See supplementary methods for a detailed listing of the parameters used). Binning of publicly available datasets was carried out using a combination of MaxBin2, Abawaca and tetranucleotide ESOM, if possible. Bins were refined using GC content, coverage and taxonomy and their completeness and contamination was accessed by a set of 51 bacterial and 38 archaeal single copy genes as described previously^11,5^. Only bins with >= 70% estimated completeness and <= 10% estimated contamination were used for downstream analysis. For each sample, genomes were dereplicated using dRep^65^.

### Ribosomal protein S3 (rpS3) analysis

Genes annotated as ribosomal protein S3 were extracted and assigned to genomes where possible based on shared GC, coverage and taxonomy. rpS3 coverage was determined based on the scaffold coverage (see above) containing the ribosomal protein. Ribosomal protein sequences were clustered using MUltiple Sequence Comparison by Log-Expectation (MUSCLE)^66^, trimmed using BMGE 1.0^67^ with the BLOSUM62 scoring matrix and aligned using IQ-TREE^68^ multicore 1.3.11.1 with -m TEST -bb^69^ 1000 and -alrt^70^ 1000 options. The tree was visualized along with other genomic data using the iToL platform version 5.5^71^.

### Determination of bacterial in situ replication indices

Reads were mapped onto concatenated genomes per sampling site using Bowtie2 with the reorder flag^60^ and the index of replication (iRep^37^) was calculated, allowing for 2% mismatches relative to the read length (3 mismatches for 150 bp). For each genome, the maximum iRep per sampling site as well as the corresponding coverage was used for downstream analysis. Coverage of each genome was determined based on the average nucleotide coverage of each scaffold across each genome (Bowtie 2) allowing also 2% mismatches. Archaea were excluded from this analysis since most of them harbor multiple origins of replication and thus the iRep signal is distorted and cannot be applied in a comparative manner. If multiple samples were available for one ecosystem, all iRep values for one genome were calculated and averaged to ensure comparability with other samples.

### Metabolic potential predictions

A set of Hidden Markov-Models (HMM) with respective score thresholds for chemolithoautotrophic key enzymes^4^ was used to predict the metabolic potential of recovered genomes and overall in entire assemblies (See suppl. Material for more detailed information).

### Biogeographical analysis

The R package sp^72^ was used to calculate the geographical elliptical distance between two sampling sites (based on longitude/latitude), in which putative genomes of the Altiarchaeales subclade Alti-1 were identified. The average nucleotide identities (ANI) between all available putative genomes of the Altiarchaeales subclade Alti-1 were calculated using the ANI calculator^73^ with default parameters. Correlations between geographical distance and ANI were done using Pearson’s r^74^.

### Genome comparison of Altiarchaeota

Genes of all Altiarchaeota genomes were blasted against each other (E-value: 10^−5^) and matches were filtered to matches with the similarity ((AlignmentLength x Identity) / QueryLength) thresholds of >= 40 %, 50 %, 60%, 70% or 80%. Cytoscape 3.7.2^75^ was used to visualize the networks at the respective similarity thresholds.

### Phylogenomic analysis of Altiarchaeota

Amino acid sequences and annotations for Alti-1 ORFs plus one Alti-2 serving as outgroup were predicted using Prokka 1.14.0^76^ with options: --kingdom archaea --metagenome --compliant). The resulting protein datasets were searched with HMMER 3.2.1^77^ for homologs of 30 universal ribosomal proteins using the v4 HMM profiles from Phylosift^78^. A 10^−4^ cutoff was applied and the resulting datasets were curated manually to remove distant homologs and multiple copies in each genome, as well as to fuse contiguous fragmented genes. Individual genes were aligned with MUSCLE v3.8.31^66^ and trimmed with BMGE^67^ under the BLOSUM30 matrix. The genes were then concatenated into a supermatrix of 5156 aa positions. The phylogeny was reconstructed in IQTree 1.6.11^68^ under the JTTDCMut+F+G4 model as selected by ModelFinder^79^.

### Community-wide analyses

Genes were predicted on assemblies with scaffolds longer than 1 kbp and chemolithoautotrophic key enzymes were predicted as described above. The abundance of the genes was estimated using the coverage of the encoding scaffolds after adjustment to unequal sequencing depths by normalization using the total bps per library. If a pathway was represented by multiple key enzymes, the enzyme with the highest frequency of hits was selected. Abundances of individual key enzymes were summed to provide the total relative abundance of each pathway in the respective samples. Likewise, diversity within each assembly was estimated based on rpS3 diversity and relative abundance of the respective scaffolds.

### Estimations of annual total erupted carbon and intracellular erupted carbon

The annual total erupted carbon was calculated based on the available CO_2_, HCO_3-_ and cell concentrations, the eruption volume (Table S1), the average estimate of the intracellular carbon amount from Whitman et al. (1998)^80^ of 90 fg cell^-1^ and the number of eruptions during tourist season (roughly 1 April – 31 October ∼ 210 days). See the Supplementary Material for the calculations.

### Statistical analysis

Statistical analyses were performed in the R programming environment^74^. These included paired and independent *t*-tests, Pearson correlations, analysis of variance (ANOVA), TukeyHSD significance tests^81^, the Shannon-Wiener index^82^ and equivalence testing using TOSTER^83^. As the upper and lower equivalence boundaries for equivalence testing of two groups, we used the effect size the CO_2_-poor sample group had a 33% power to detect as recommended previously^84^. Results were visualized using ggplot2^85^.

***Methods for DAPI staining, cell counting, geochemical measurements are provided in the Supplementary Methods***

## Acknowledgements

This study was funded by the Ministerium für Kultur und Wissenschaft des Landes Nordrhein-Westfalen (“Nachwuchsgruppe Dr. Alexander Probst”). We thank Hubert Müller for technical assistance and Karen L Lloyd for scientific discussions.

## Author contribution

TLVB performed main bioinformatics analysis. PSA performed phylogenomics. VT and AJP performed microscopy. US, RS and BK performed geological analyses and geological data interpretation. TLVB and PAFG analyzed genomes. TLVB, JR and AJP took samples. DK and TCS performed geochemical analyses. AJP conceptualized the study. TLVB and AJP wrote the manuscript with revisions from all co-authors.

## Notes

#### Summary of Updates

Additional analyses were added to the supplementary. Main figures were visually improved.

## References

1. Magnabosco, C. et al. The biomass and biodiversity of the continental subsurface. Nat. Geosci. 11, 707–717 (2018).

2. Flemming, H.-C. & Wuertz, S. Bacteria and archaea on Earth and their abundance in biofilms. Nat. Rev. Microbiol. 17, 247–260 (2019).

3. Falkowski, P. G., Fenchel, T. & Delong, E. F. The microbial engines that drive Earth’s biogeochemical cycles. Science 320, 1034–1039 (2008).

4. Anantharaman, K. et al. Thousands of microbial genomes shed light on interconnected biogeochemical processes in an aquifer system. Nat. Commun. 7, 1–11 (2016).

5. Probst, A. J. et al. Differential depth distribution of microbial function and putative symbionts through sediment-hosted aquifers in the deep terrestrial subsurface. Nat. Microbiol. 3, 328–336 (2018).

6. Castelle, C. J. et al. Genomic Expansion of Domain Archaea Highlights Roles for Organisms from New Phyla in Anaerobic Carbon Cycling. Curr. Biol. 25, 690–701 (2015).

7. Stevens, T. Lithoautotrophy in the subsurface. FEMS Microbiol. Rev. 20, 327–337 (1997).

8. Stevens, T. O. & McKinley, J. P. Abiotic Controls on H_2_ Production from Basalt−Water Reactions and Implications for Aquifer Biogeochemistry. Environ. Sci. Technol. 34, 826–831 (2000).

9. Nyyssönen, M. et al. Taxonomically and functionally diverse microbial communities in deep crystalline rocks of the Fennoscandian shield. ISME J. 8, 126–138 (2014).

10. Lau, M. C. Y. et al. An oligotrophic deep-subsurface community dependent on syntrophy is dominated by sulfur-driven autotrophic denitrifiers. Proc. Natl. Acad. Sci. 113, E7927–E7936 (2016).

11. Probst, A. J. et al. Genomic resolution of a cold subsurface aquifer community provides metabolic insights for novel microbes adapted to high CO_2_ concentrations. Environ. Microbiol. 19, 459–474 (2017).

12. Probst, A. J. et al. Biology of a widespread uncultivated archaeon that contributes to carbon fixation in the subsurface. Nat. Commun. 5, 5497 (2014).

13. Hernsdorf, A. W. et al. Potential for microbial H_2_ and metal transformations associated with novel bacteria and archaea in deep terrestrial subsurface sediments. ISME J. 11, 1915–1929 (2017).

14. Bird, J. T., Baker, B. J., Probst, A. J., Podar, M. & Lloyd, K. G. Culture Independent Genomic Comparisons Reveal Environmental Adaptations for Altiarchaeales. Front. Microbiol. 7, (2016).

15. Moissl, C., Rachel, R., Briegel, A., Engelhardt, H. & Huber, R. The unique structure of archaeal ‘hami’, highly complex cell appendages with nano-grappling hooks: Unique structure of archaeal ‘hami’. Mol. Microbiol. 56, 361–370 (2005).

16. Wood, H. G. Life with CO or CO_2_ and H_2_ as a source of carbon and energy. FASEB J. 5, 156–163 (1991).

17. Gutiérrez-Preciado, A. et al. Functional shifts in microbial mats recapitulate early Earth metabolic transitions. Nat. Ecol. Evol. 2, 1700–1708 (2018).

18. Adam, P. S., Borrel, G. & Gribaldo, S. An archaeal origin of the Wood–Ljungdahl H 4 MPT branch and the emergence of bacterial methylotrophy. Nat. Microbiol. 4, 2155–2163 (2019).

19. Bräuer, K., Kämpf, H., Niedermann, S. & Strauch, G. Indications for the existence of different magmatic reservoirs beneath the Eifel area (Germany): A multi-isotope (C, N, He, Ne, Ar) approach. Chem. Geol. 356, 193–208 (2013).

20. Zhang, Y. Degassing History of Earth. in Treatise on Geochemistry 37–69 (2014).

21. Caracausi, A. & Paternoster, M. Radiogenic helium degassing and rock fracturing: A case study of the southern Apennines active tectonic region. J. Geophys. Res. Solid Earth 120, 2200–2211 (2015).

22. Loreto, M. F., Italiano, F., Deponte, D., Facchin, L. & Zgur, F. Mantle degassing on a near shore volcano, SE Tyrrhenian Sea. Terra Nova 27, 195–205 (2015).

23. Gilfillan, S. M. V. et al. Noble gases confirm plume-related mantle degassing beneath Southern Africa. Nat. Commun. 10, 1–7 (2019).

24. Lee, H. et al. Mantle degassing along strike-slip faults in the Southeastern Korean Peninsula. Sci. Rep. 9, 1–9 (2019).

25. Hedrick, D. B., Pledger, R. D., White, D. C. & Baross, J. A. In situ microbial ecology of hydrothermal vent sediments. FEMS Microbiol. Lett. 101, 1–10 (1992).

26. Schrenk, M. O., Holden, J. F. & Baross, J. A. Magma-to-microbe networks in the context of sulfide hosted microbial ecosystems. Wash. DC Am. Geophys. Union Geophys. Monogr. Ser. 178, 233–258 (2008).

27. Ding, J. et al. Microbial Community Structure of Deep-sea Hydrothermal Vents on the Ultraslow Spreading Southwest Indian Ridge. Front. Microbiol. 8, (2017).

28. Tu, T.-H. et al. Microbial Community Composition and Functional Capacity in a Terrestrial Ferruginous, Sulfate-Depleted Mud Volcano. Front. Microbiol. 8, (2017).

29. Galambos, D., Anderson, R. E., Reveillaud, J. & Huber, J. A. Genome-resolved metagenomics and metatranscriptomics reveal niche differentiation in functionally redundant microbial communities at deep-sea hydrothermal vents. Environ. Microbiol. 21, 4395–4410 (2019).

30. Frerichs, J. et al. Microbial community changes at a terrestrial volcanic CO_2_ vent induced by soil acidification and anaerobic microhabitats within the soil column. FEMS Microbiol. Ecol. 84, 60–74 (2013).

31. Mehlhorn, J., Beulig, F., Küsel, K. & Planer-Friedrich, B. Carbon dioxide triggered metal(loid) mobilisation in a mofette. Chem. Geol. 382, 54–66 (2014).

32. Beulig, F. et al. Carbon flow from volcanic CO_2_ into soil microbial communities of a wetland mofette. ISME J. 9, 746–759 (2015).

33. Beulig, F. et al. Altered carbon turnover processes and microbiomes in soils under long-term extremely high CO_2_ exposure. Nat. Microbiol. 1, 1–10 (2016).

34. Mori, K., Yamaguchi, K., Sakiyama, Y., Urabe, T. & Suzuki, K. Caldisericum exile gen. nov., sp. nov., an anaerobic, thermophilic, filamentous bacterium of a novel bacterial phylum, Caldiserica phyl. nov., originally called the candidate phylum OP5, and description of Caldisericaceae fam. nov., Caldisericales ord. nov. and Caldisericia classis nov. Int. J. Syst. Evol. Microbiol. 59, 2894–2898 (2009).

35. Sharrar, A. M. et al. Novel Large Sulfur Bacteria in the Metagenomes of Groundwater-Fed Chemosynthetic Microbial Mats in the Lake Huron Basin. Front. Microbiol. 8, (2017).

36. Hoang, D. T., Chernomor, O., von Haeseler, A., Minh, B. Q. & Vinh, L. S. UFBoot2: Improving the Ultrafast Bootstrap Approximation. Mol. Biol. Evol. 35, 518–522 (2018).

37. Guindon, S. et al. New algorithms and methods to estimate maximum-likelihood phylogenies: assessing the performance of PhyML 3.0. Syst. Biol. 59, 307–321 (2010).

38. Anisimova, M., Gil, M., Dufayard, J.-F., Dessimoz, C. & Gascuel, O. Survey of Branch Support Methods Demonstrates Accuracy, Power, and Robustness of Fast Likelihood-based Approximation Schemes. Syst. Biol. 60, 685–699 (2011).

39. Kairesalo, T., Tuominen, L., Hartikainen, H. & Rankinen, K. The role of bacteria in the nutrient exchange between sediment and water in a flow-through system. Microb. Ecol. 29, 129–144 (1995).

40. Brown, C. T., Olm, M. R., Thomas, B. C. & Banfield, J. F. Measurement of bacterial replication rates in microbial communities. Nat. Biotechnol. 34, 1256 (2016).

41. Zerulla, K. et al. DNA as a Phosphate Storage Polymer and the Alternative Advantages of Polyploidy for Growth or Survival. PLoS ONE 9, (2014).

42. Frawley, L. E. & Orr-Weaver, T. L. Polyploidy. Curr. Biol. 25, R353–R358 (2015).

43. Lennon, J. T. & Jones, S. E. Microbial seed banks: the ecological and evolutionary implications of dormancy. Nat. Rev. Microbiol. 9, 119–130 (2011).

44. Locey, K. J., Fisk, M. C. & Lennon, J. T. Microscale Insight into Microbial Seed Banks. Front. Microbiol. 7, (2017).

45. Jorgensen, S. L. et al. Correlating microbial community profiles with geochemical data in highly stratified sediments from the Arctic Mid-Ocean Ridge. Proc. Natl. Acad. Sci. U. S. A. 109, E2846–E2855 (2012).

46. Wright, S. Isolation by Distance. Genetics 28, 114–138 (1943).

47. Wright, S. Isolation by Distance Under Diverse Systems of Mating. Genetics 31, 39–59 (1946).

48. Malecot, G. Mathematics of heredity. Math. Hered. (1948).

49. Martiny, J. B. H. et al. Microbial biogeography: putting microorganisms on the map. Nat. Rev. Microbiol. 4, 102–112 (2006).

50. Schunk, R. Der Ausbruch – ein faszinierendes Naturschauspiel. in Naturschauspiel Geysir Andernach 20–36 (2012).

51. Krauthausen, B., Deuster, J. & Lang, R. Die Flucht des Wassers aus der Tiefe. Der Geysir von Andernach am Rhein. in Faszination Geologie. Die bedeutendsten Geotope Deutschlands 110–111 (2007).

52. Altfeld, E. Die physikalischen Grundlagen des intermittierenden Kohlensäuresprudels zu Namedy bei Andernach a. Rh. (1913).

53. Dèzes, P., Schmid, S. M. & Ziegler, P. A. Evolution of the European Cenozoic Rift System: interaction of the Alpine and Pyrenean orogens with their foreland lithosphere. Tectonophysics 389, 1–33 (2004).

54. Meyer, W. & Stets, J. Geologische Übersichtskarte und Profil des Mittelrheintales - 1:100000, mit Erläuterungen. in 49 ((Geologisches Landesamt Rheinland-Pfalz) Mainz, 2000).

55. Meyer, W. & Striem, H. L. Geological indications for young horizontal displacements in the Central Rhenish Massif. Geol. Indic. Young Horiz. Displac. Cent. Rhenish Massif 97–100 (1983).

56. Schreiber, U. & Rotsch, S. Cenozoic block rotation according to a conjugate shear system in central Europe — indications from palaeomagnetic measurements. Tectonophysics 299, 111–142 (1998).

57. Ritter, J. R. R. The Seismic Signature of the Eifel Plume. Mantle Plumes, 379–404 (2007).

58. JN Fass, N. J. Sickle: A sliding-window, adaptive, quality-based trimming tool for FastQ files. (2011).

59. Nurk, S., Meleshko, D., Korobeynikov, A. & Pevzner, P. A. metaSPAdes: a new versatile metagenomic assembler. Genome Res. 27, 824–834 (2017).

60. Hyatt, D. et al. Prodigal: prokaryotic gene recognition and translation initiation site identification. BMC Bioinformatics 11, 119 (2010).

61. Buchfink, B., Xie, C. & Huson, D. H. Fast and sensitive protein alignment using DIAMOND. Nat. Methods 12, 59–60 (2015).

62. Suzek, B. E., Huang, H., McGarvey, P., Mazumder, R. & Wu, C. H. UniRef: comprehensive and non-redundant UniProt reference clusters. Bioinformatics 23, 1282–1288 (2007).

63. Langmead, B. & Salzberg, S. L. Fast gapped-read alignment with Bowtie 2. Nat. Methods 9, 357–359 (2012).

64. Wu, Y.-W., Simmons, B. A. & Singer, S. W. MaxBin 2.0: an automated binning algorithm to recover genomes from multiple metagenomic datasets. Bioinformatics 32, 605–607 (2016).

65. Dick, G. J. et al. Community-wide analysis of microbial genome sequence signatures. Genome Biol. 10, R85 (2009).

66. Alneberg, J. et al. Binning metagenomic contigs by coverage and composition. Nat. Methods 11, 1144–1146 (2014).

67. Sieber, C. M. K. et al. Recovery of genomes from metagenomes via a dereplication, aggregation and scoring strategy. Nat. Microbiol. 3, 836–843 (2018).

68. Olm, M. R., Brown, C. T., Brooks, B. & Banfield, J. F. dRep: a tool for fast and accurate genomic comparisons that enables improved genome recovery from metagenomes through de-replication. ISME J. 11, 2864 (2017).

69. Edgar, R. C. MUSCLE: multiple sequence alignment with high accuracy and high throughput. Nucleic Acids Res. 32, 1792–1797 (2004).

70. Criscuolo, A. & Gribaldo, S. BMGE (Block Mapping and Gathering with Entropy): a new software for selection of phylogenetic informative regions from multiple sequence alignments. BMC Evol. Biol. 10, 210 (2010).

71. Nguyen, L.-T., Schmidt, H. A., von Haeseler, A. & Minh, B. Q. IQ-TREE: A Fast and Effective Stochastic Algorithm for Estimating Maximum-Likelihood Phylogenies. Mol. Biol. Evol. 32, 268–274 (2015).

72. Letunic, I. & Bork, P. Interactive tree of life (iTOL) v3: an online tool for the display and annotation of phylogenetic and other trees. Nucleic Acids Res. 44, W242–W245 (2016).

73. Pebesma, E. & Bivand, R. Classes and Methods for Spatial Data in R. R News 5, (2005).

74. Rodriguez-R, L. M. & Konstantinidis, K. T. The enveomics collection: a toolbox for specialized analyses of microbial genomes and metagenomes. https://peerj.com/preprints/1900 (2016)

75. R Core Team. R: A Language and Environment for Statistical Computing. R Foundation for Statistical Computing, Vienna, Austria (2008).

76. Shannon, P. et al. Cytoscape: a software environment for integrated models of biomolecular interaction networks. Genome Res. 13, 2498–2504 (2003).

77. Seemann, T. Prokka: rapid prokaryotic genome annotation. Bioinforma. Oxf. Engl. 30, 2068–2069 (2014).

78. Eddy, S. R. Accelerated Profile HMM Searches. PLoS Comput. Biol. 7, e1002195 (2011).

79. Darling, A. E. et al. PhyloSift: phylogenetic analysis of genomes and metagenomes. PeerJ 2, e243 (2014).

80. Kalyaanamoorthy, S., Minh, B. Q., Wong, T. K. F., von Haeseler, A. & Jermiin, L. S. ModelFinder: fast model selection for accurate phylogenetic estimates. Nat. Methods 14, 587–589 (2017).

81. Whitman, W. B., Coleman, D. C. & Wiebe, W. J. Prokaryotes: The unseen majority. Proc. Natl. Acad. Sci. 95, 6578–6583 (1998).

82. Haynes, W. Tukey’s Test. in Encyclopedia of Systems Biology (eds. Dubitzky, W., Wolkenhauer, O., Cho, K.-H. & Yokota, H.) 2303–2304 (Springer New York, 2013).

83. Shannon, C. E. A mathematical theory of communication. Bell Syst. Tech. J. 27, 379–423 (1948).

84. Lakens, D., Scheel, A. M. & Isager, P. M. Equivalence Testing for Psychological Research: A Tutorial: Adv. Methods Pract. Psychol. Sci. (2018).

85. Simonsohn, U. Small Telescopes: Detectability and the Evaluation of Replication Results. Psychol. Sci. 26, 559–69 (2015).

86. Wickham, H. ggplot2: Elegant Graphics for Data Analysis. (Springer-Verlag, 2009).

