## Supplementary Information for "Geological degassing enhances microbial metabolism in the continental subsurface"

Main Supplementary File for:

**Content**:

1. Supplementary Methods
2. Supplementary Discussions
3. Supplementary Figures
4. Supplementary Tables
5. Additional supplementary files
6. References

### SUPPLEMENTARY METHODS

***Estimations of annual total erupted carbon and intracellular erupted carbon.*** The annual total erupted carbon was calculated based on the available $CO_2$ and $HCO_3$ concentrations and the eruption volume (Table S1) and the average estimate of the intracellular carbon amount from Whitman et al. (1998)[1] of 90 fg/cell. During tourist season (April 1 – October 31, ~ 210 days, rest of the year no eruptions), three eruptions are initiated per day with an average eruption volume of 6.5 $m^3$. We used equation Eq (1) to calculate the total erupted organic carbon and Eq (2) to calculate the total erupted cellular carbon.

Eq (1) $Total\ carbon = V\ x\ E\ x\ D\ x\ (CO2xFCO2 + HCO3\ x\ FHCO3) = 6.27\ t\ carbon/year$

$Eq\ (2) Cellular\ carbon = Vx\ E\ x\ D\ x\ C\ x\ cellular\ carbon\ =\ 716.625\ g$

| **Abbreviations** | **Description** | **Value** |
|---|---|---|
| V | eruption volume | 6.5 $m^3$ |
| E | number of eruptions per day | 3 |
| D | tourist season length in days | 210 (7 months) |
| $CO_2$ | concentration of $HCO_3$ | 1500 mg / l |
| $F_{CO2}$ | mass fraction of C in $CO_2$ | 0.27 |
| $HCO_3$ | concentration of $HCO_3$ | 5700 mg / l |
| $F_{HCO3}$ | mass fraction of C in $HCO_3$ | 0.20 |
| C | cell concentration | 3.5 x $10^6$ cells $ml^{-1}$ |

***Phylogenetic placement of draft genomes.*** Marker genes *rpL16*, *rpL18*, *rpL2*, *rpL22*, *rpL24*, *rpL3*, *rpL4*, *rpL5*, *rpL6*, *rpS10*, *rpS17*, *rpS19*, *rpS3* and *rpS8* were identified in the draft genomes using blastp with an e-value cutoff of 0.01 against a previously described database of bacterial homologues to these marker genes[2] and verified by blasting of the resulting candidate sequences against all proteins of all genomes in the database with an e-value cutoff of $10^{-6}$.

***Metagenomic binning.*** Scaffolds of each sample were binned using ABAWACA and ESOMs [3,4] based on tetra-nucleotide frequency of DNA fragments of 3 kbp and 5 kbp as minimum length and 5 kbp and 10 kbp as respective maximum length cutoffs, respectively. *Escherichia coli* (low GC%) and *Streptomyces griseus* (high GC%) spike-in genomes prior to nucleotide frequency calculation were used to verify the success of the ESOM training and only ESOMs showing a good clustering of these controls were used for binning. SulCav samples were not binned using ESOM due to sample size. Differential coverage binning was achieved using Maxbin2[5] (default parameters), if multiple samples were available for the same ecosystem. GA samples were additionally binned using CONCOCT[6] (default parameters). The different binning results were aggregated using DAS Tool[7] (default parameters) and the genomic bins were curated based on their GC content, coverage and taxonomy. Completeness and contamination was assessed using a previously described set of universal 51 bacterial and 38 archaeal single copy genes[8] and only genomes with >= 70 % completeness and <= 10 % contamination were used for further analyses. Genomes originating from different samples of the same ecosystem were dereplicated using dRep[9].

***Geochemical measurements of metagenomic samples*.**

*Temperature.* The air temperature was measured using a mobile phone and the water temperature using thermometers in triplicate.

*pH.* The pH was measured using pH strips with 0.2 Interval coloring schemes in triplicate.
*Sulfide.* 200 ul of water were added to 1 ml of 1 % (w/v) Zinc-Acetate and mixed on-site. Concentrations were measured according to a modified version of the protocol by Cline[10] in triplicate. 20 µl of sample was added to 400 µl of 1 % (w/v) Zinc-Acetate and 0.2 % (v/v) acetic acid. 25 µl of both 0.5 % (w/v) ferric ammonium sulphate + 0.096 % (v/v) sulfuric acid and of 0.2 % (w/v) 4-amino-N,N-dimethylaniline sulphate + 19.6 % (v/v) sulfuric acid were added to each well of the plate reader, followed by addition of 100 µl of diluted sample. The plate was incubated in the dark for 30 minutes followed by detection of absorbance at 664 nm. The sulfide concentration was determined using a dilution series of 0.05 mM to 2 mM of sulfide. Measurements were performed in triplicate and triplicates were averaged.
*Fe(II) determination.* 100 ul of sample were added to 900 ul of 1 M HCl and mixed on site. Fe(II) concentration was determined by addition of 20 µl of sample to 180 µl of 0.1 % (w/v) ferrozine with 50 % (w/v) ammonium acetate in plate reader plate wells, incubation in dark for 15 minutes followed by absorbance detection at 560 nm. A dilution series from 0 mM to 100 mM was used to determine the concentration.
*Total iron determination.* 100 ul of sample were added to 900 ul of 1 M HCl and mixed on site. 100 µl of the mixture (as well as the dilution series of Fe(II) concentration determination) were added to 900 µl of 10 % (w/v) hydroxylamine-HCl in 1 M HCl and incubated on a shaker for 15 minutes to dissolve iron precipitates. Then 20 µl of solution are added to 180 µl of 0.1 % (w/v) ferrozine with 50 % (w/v) ammonium acetate in the microtiter plate and the absorbance is detected at 560 nm.
*Total organic carbon (TOC) measurements.* 2 ml of 1 M HCl were added to 10 ml of filtered sample (0.45 um pore size) and degassed by bubbling air into the vials for 15 min to purge the sample of inorganic carbon. The TOC concentrations were determined using a TOC-L (Shimadzu). Two biological replicates were measured with three technical replicates.
*Ion measurements.* 0.1 um pore-size filtrated water was used for ion measurements. Samples were measured in dilutions of 1:100 and 1:600 in distilled water to be able to accurately quantify both highly abundant ions and lower abundant ions. Two biological samples were analyzed using the Dionex Aquion ion chromatography system (Thermo Scientific, USA). Anions were analyzed with a Dionex IonPac AG23-4 µm guard column, a Dionex IonPac AG23-4 µm 2 x 250 mm analytical column as well as an AERS 500 Carbonate 2 mm suppressor, Ultimate 3000 heating element and a DS6 heated conductivity cell detector. Cations were measured on a CS12A RFIC 2x 250 mm analytical column and a CG12A RFIC detector. Measurements were done in technical triplicates and distilled water blank concentrations were subtracted.

***Long-term geochemical measurements*** were performed according to German TrinwV-GW (drinking water guidelines).

***DAPI Staining and cell enumeration.***
*Staining.* Water was filtered on-site through 0.1 um pore size filters. Filters were incubated with 10 ug/ml DAPI in 2 % (v/v) Formaldehyde for 5 minutes, washed with 30 ml of distilled water, dried for 10 minutes and stored in the dark at 4° C till use. All steps involving DAPI and stained filters were done in the dark.
*Enumeration of cells.* Cells were quantified in cells $ml^{-1}$ using the Axio A.1 epiflouorescence Microscope at by enumeration of the cells in ten horizontal and ten vertical fields, extrapolating counts to the entire filter area and normalizing through the filtered volume.

### SUPPLEMENTARY DISCUSSION

#### Geography and dispersal of Altiarchaeota

All Alti-1 species (including two new bins from GA and SulCav) placed the under the Altiarchaota subclade Alti-1 based on our phylogenomic analysis. Alti-1 organisms are known to dominate their respective subsurface ecosystems and form biofilms using their characteristic hook-like *hami* surface appendages[11,12]. Our phylogenomic analysis reproduced prior findings by Bird[13] and expanded upon the Alti-1 subclade phylogenetic diversity. Based on our analysis, Alti-1 genomes have an extreme degree of provincialism, with clear clustering by continent of origin (North America, Europe, Asia), which is also reproducible based on ANI and AAI as estimates of genome similarity. Prior intercontinental studies have also observed a continental clustering for their respective *Synechococcus*[14], *Sulfolobus*[15] or *Comamonas testosteroni*[16] assemblages and identified geographical distance as the main predictor as opposed to other environmental parameters[14,15].

The relationship between genomic similarity and geographic distance is in general a well-known occurrence in nature and as a consequence, concepts like the isolation-by-distance theory[17,18,19] and others[20,21] have been developed. Generally, isolation-by-distance suggests that distance to be the main barrier for dispersal and consequently limiting the exchange of genetic material. Such relationships are also well-established for microorganisms, with atmospheric and hydrological phenomena contributing to the dispersal[22,23]. However, microorganisms hitchhiking on plate tectonics for their dispersal is not unheard of either[24,25]. Since Altiarchaeales are anaerobic subsurface dwellers, surficial dispersal mechanisms or subsurface dispersal via oxygenated groundwater systems are unlikely to have contributed to their spread across multiple continents. Their *hami*, have been shown to be very adhesive to a variety of different surfaces, in addition to being rather heat- and pH-stable, thus likely anchoring Altiarcheales biofilms in their location[11]. The absence of evidence for *hami* is also one of the major differences to the subclade Alti-2, which is much more genetically diverse and exhibits more complex biogeographical patterns[13] (see File S2), potentially indicating a larger degree of dispersal. This makes plate tectonics the most likely dispersal route for organisms of the Alti-1 clade. Even within the Phanerozoic, there were ample opportunities for the common ancestor of Alti1 to result in the North American and European clades, starting with the early Devonian (~400 Ma), when the two continental margins Laurentia and Baltica (making up North America and Europe, respectively) collided to form Laurasia[26,27]. Japan on the other hand has not been in direct contact with Europe or North America since the break-up of the supercontinent Rodinia 750-600 Ma years ago[28] and thus there are not really any opportunities for direct dispersal between the other sampling sites and the Japanese site during the Phanerozoic. The European and Japanese Alti-1 clades show closer similarity between themselves than to the American clade, indicating that they share a common ancestor. A potential dispersal route for their ancestral populations could be across the Siberian plate through China by the early Mesozoic and then transferred to Japan during the tectonic processes, by which it emerged from the sea 25 Ma ago. However, as there are currently no Altiarchaeales Alti-1 genomes available from these possible intermediate transfer points as well as possible negative controls that are outside of this path of dispersal, this remains speculative.

**Geological reasoning behind the enhancement of microbial activity in ecosystems affected by geological degassing**

In our study, we observed that microbes in the Geyser Andernach and in the Crystal Geyser replicated much faster than they should, based on the observed correlation between bacterial replication indices as estimated using iRep and the sampling depth and we proposed that the higher nutrient flow rates present in these ecosystems could very well explain this difference. In the following, the geological reasoning behind the proposal will be explained in more detail.

In sediments, aquifers flow through pore channels in the subsurface and their amount generally decreases with depth as the sediment becomes more compact and the pores mineralize, thus reducing the diameter and consequently the flow rate. Joint aquifers, which are characterized by solid rock formation-embedded channels, can allow for much greater flowrates that can be up to multiple magnitudes greater than sediment pore channel aquifers since no compression occurs. In aquifers, the nutrient availability is directly proportional to the flow volume of the groundwater and thus the greater flow rate in joint aquifers should automatically increase the microbial activity since nutrient availability is the generally the growth limiting factor in the subsurface[29].

Ecosystems like the Geyser Andernach or mofettes that are affected by mantle degassing are such joint aquifer systems since the release of the $CO_2$-rich gases requires complex channel systems with a mixture of water and $CO_2$ with traces of e.g. hydrogen, $SH_2$, methane and nitrogen. Within the mantle, organic products are generated by the combination of high pressure and temperature comparable to the Fischer/Tropsch-Synthesis. These products are mainly long-chain alkanes depending on the depth, gas composition and availability of metals on crust surfaces. Below 20 km depth, both water and $CO_2$ are supercritical and infinitely mixable. Decreasing pressure liquifies the water, segregating the $CO_2$ and causing it to rise to the surface as droplets, capturing the nonpolar alkanes in its current. Once $CO_2$ becomes subcritical, the organics precipitate at the interface since the gaseous $CO_2$ cannot keep them in solution. Gaseous $CO_2$ then travels on to the surface and initiates a current, transporting the organics. This current results in a steady supply of organics as well as gases like $CO_2$, $H_2$ and $N_2$, fueling microbial activity and replication and is thus a probable explanation for the comparatively high replication measures in high $CO_2$ environments.

**SUPPLEMENATARY FIGURES**

**Fig. S1 | Cell densities in Geyser Andernach in comparison to other sample types and depths.** Cell concentrations across different sample types and depths from Magnabosco[30] (colored dots) are set in relation to the concentration of the Geyser Andernach (black dot, mean cell concentration is shown). To increase visibility of the Geyser Andernach sample, only cell concentrations of ecosystems with 0-500 m depth and with relevant sample types are shown.

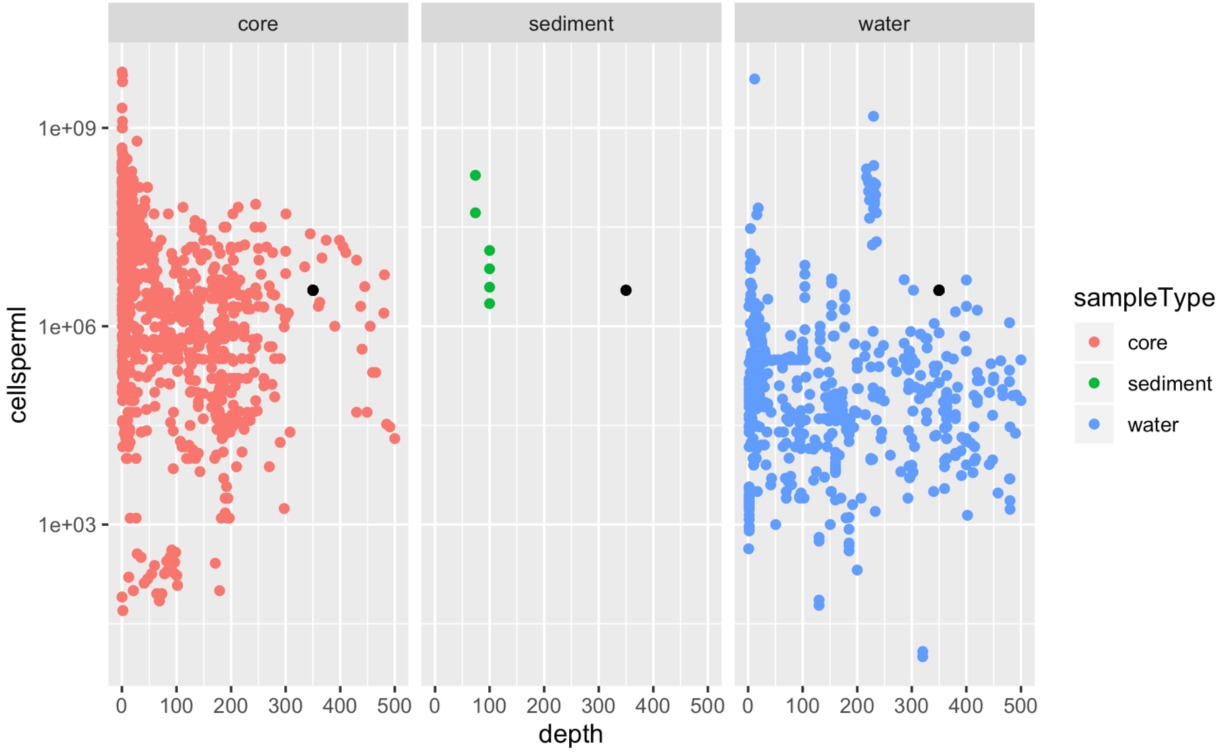

**Fig. S2 | Correlation of ANI (A) and AAI (B) similarity of Alti-1 genomes with geographical distance. A:** Correlation of average nucleotide identity (ANI) in dependency on the geographical distance between sampling sites. The elliptical geographical distance (as opposed to the Euclidian distance) was determined using the R package sp[31] and the ANI was determined using fastANI[32]. Pearson correlation was used to see whether the two variables show a dependency. **B:** Correlation of average amino acid identity (AAI) in dependency on the geographical distance between sampling sites. The elliptical geographical distance (as opposed to the Euclidian distance) was determined using the R package sp and the AAI was determined using fastANI. Pearson correlation was used to see whether the two variables show a dependency.

**A**

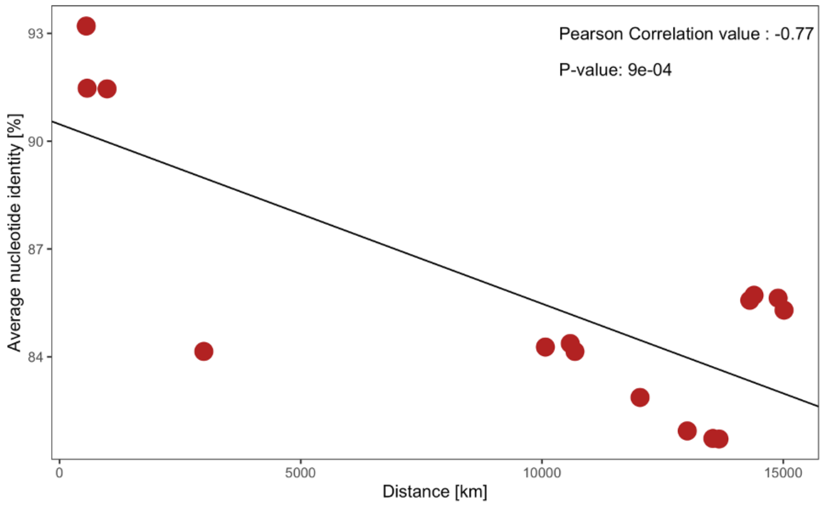


**B**

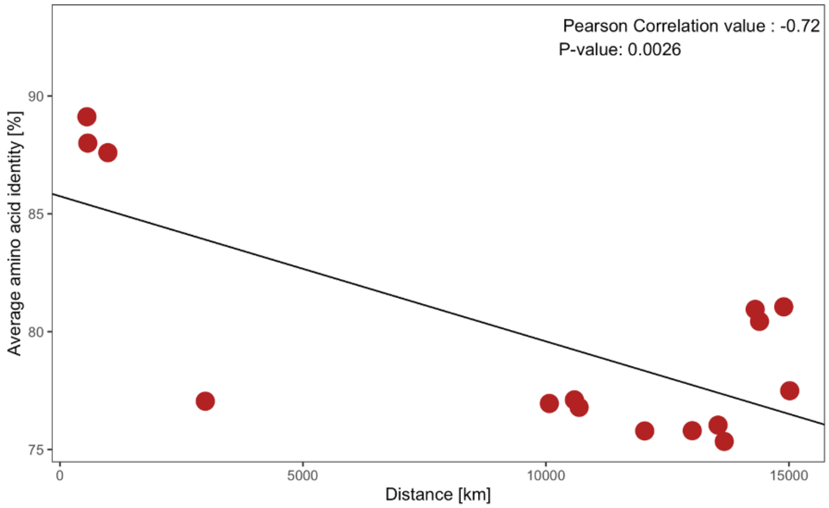

**Fig. S3 | Phylogeny of Alti-1 genotypes based on 30 universal ribosomal proteins** (5136 aa positions, IQTree JTTDCMut+F+G4) and using the Alti-2 genome IMC4 as the outgroup. Branch supports correspond to ultrafast bootstraps[33] (1000 replicates), the SH-aLRT[34] test (1000 replicates), and the approximate Bayes test[35] respectively.

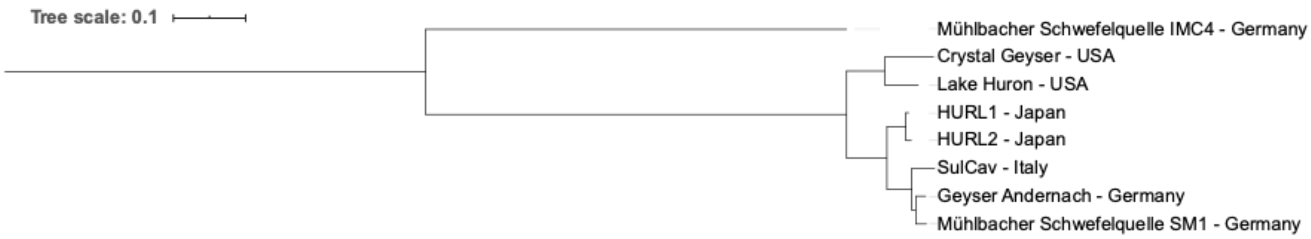


**Fig. S4 | Correlation between Shannon-index and depth.** Shannon indices were calculated using the R package vegan[36] from sequencing depth-normalized rpS3-scaffold abundances for each sample. The median for each sampling depth was computed and then correlated against the sampling site depth using Pearson correlation.

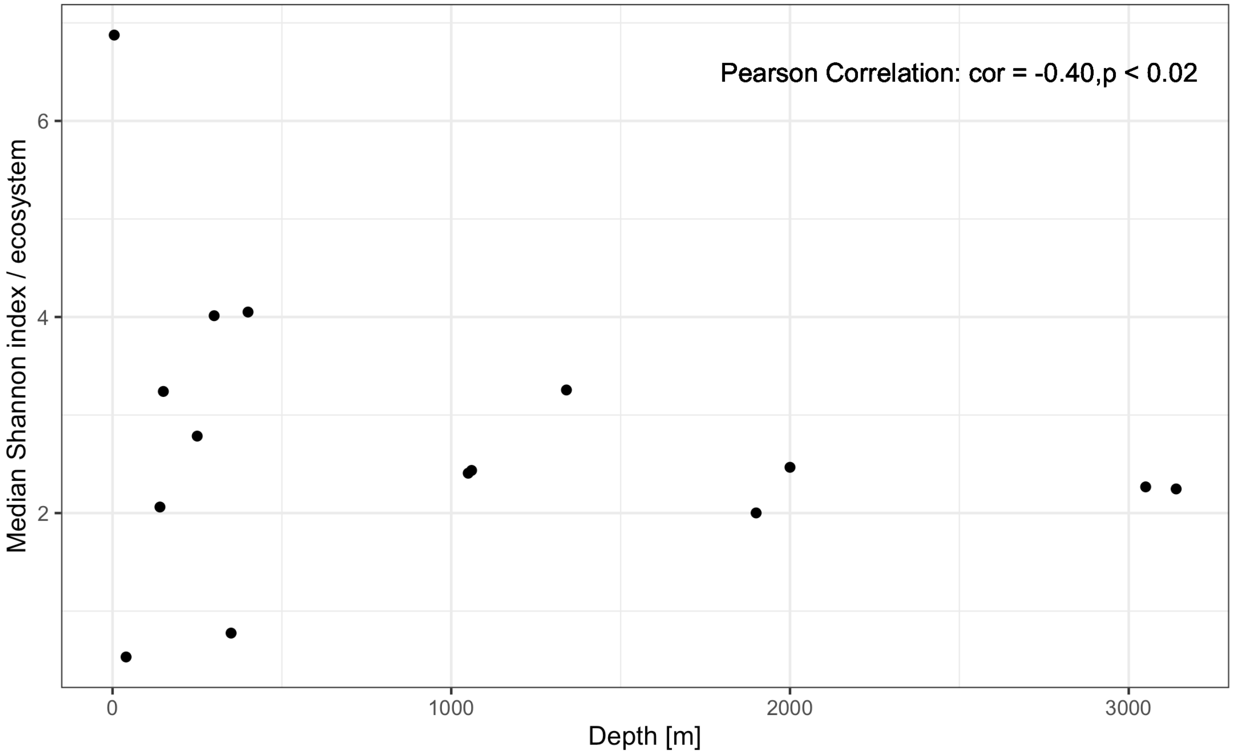

**Figure S5 | Protein similarity clustering networks of Altiarchaeales proteins with >= 80% similarity. A:** Genomes are marked as large red circles with edges marked as black lines. Edge lengths are arbitrary. Genomes belonging to the Alti-1 subclade are enclosed by a square. The network was visualized in Cytoscape[37]. **B:** Presence / absence heatmap of protein clusters (y-axis) across Altiarchaeales genomes (x-Axis) with an 80 % similarity cutoff and a hierarchical clustering dendrogram showing the overall relatedness of Altiarchaeles proteins.

**A** **B**

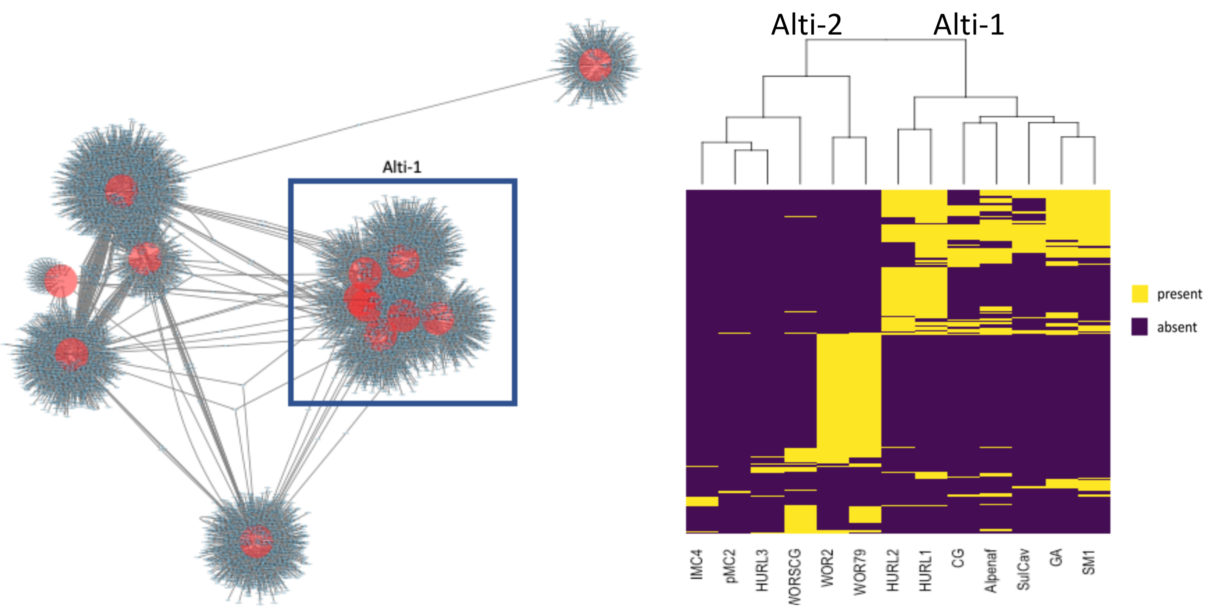

### SUPPLEMENTARY TABLES

**Table S1 | Geochemical measurements of Geyser Andernach.** Additional data can be found in additional file 3.

**General characterization**

| | |
|---|---|
| Eruption interval | 120 min |
| Eruption duration | 6 (2004) – 30 (2018) min |
| Maximum eruption height | 45 - 62 m |
| Eruption water volume | 6 - 7m³ / eruption |
| Temperature water at surface | 18 - 22° C |
| Water temperature in 240 m depth | 26.5 °C |
| $CO_2$ temperature at surface | 20 - 22° C |
| Water Conductivity | 12.800 - 13.600 µS |
| *potentia hydrogenii* (pH) | 6.6 – 6.8 |
| Chemical composition (averages) | |
| Sodium [$Na^+$] | 2.750 mg $l^{-1}$ |
| Kalium [$K^+$] | 100 mg $l^{-1}$ |
| Magnesium [$Mg^{2+}$] | 340 mg $l^{-1}$ |
| Calcium [$Ca^{2+}$] | 300 mg $l^{-1}$ |
| Manganese [Mn] | <1 mg $l^{-1}$ |
| Iron [Fe] | 10 mg $l^{-1}$ |
| Chloride [$Cl^-$] | 2.250 mg $l^{-1}$ |
| Sulfate [$SO_4^{2-}$] | 425 mg $l^{-1}$ |
| Hydrogen carbonate [$HCO_3^-$] | 5.700 mg $l^{-1}$ |
| Carbon dioxide [$CO_2$] | 1.500 mg $l^{-1}$ |

**Gas composition**[38]

| | |
|---|---|
| Carbon dioxide [$CO_2$] | 99.96 % (v/v) |
| Hydrogen [$H_2$] | 2.4 ppmv |
| Oxygen [$O_2$] | 0.008 % (v/v) |
| Nitrogen [$N_2$] | 0.03 % (v/v) |
| Methane [$CH_4$] | BDL |
| Helium [He] | 1.6 ppmv |
| Argon [Ar] | 0.0007 % (v/v) |

**Chemical analysis of samples also investigated with metagenomics**

| Compound (unit) | AVG (STDEV) |
|---|---|
| Calcium $Ca^{2+}$(mM) | 5.26 (0.21) |
| Magnesium $Mg^{2+}$ (mM) | 9.05 (0.27) |
| Potassium $K^{+}$ (mM) | 3.15 (0.86) |
| Ammonia $NH_4^{+}$ (mM) | 1.20 (0.35) |
| Sodium $Na^{+}$ (mM) | 73.05 (2.28) |
| Sulfate $SO_4^{2-}$ (mM) | 4.98 (0.39 |
| Chloride $Cl^{-}$ (mM) | 70.58 (7.26) |
| Nitrate $NO_3^{-}$ (mM) | BDL |
| Phosphate $PO_4^{3-}$ (mM) | BDL |
| ferrous iron $Fe^{2+}$ (mM) | BDL |
| $Fe_{total}$ (μM) | 486.11 (39.06) |
| $H_2S$ (μM) | BDL |
| DOC (mg / l) | 1.39 (0.31) |

**Table S2 | Assembly statistics** (all samples, all ecosystems).

| Assembly | Publication of read dataset | # TB read bps | % reads mapped to assembly | # scaffold bps | # scaffolds >=1000 bps length | N50 scaffolds >=1000 bps length |
|---|---|---|---|---|---|---|
| GA_E1_1 | This study | 6.9 | 75 | 3.66E+08 | 1.08E+08 | 3090 |
| GA_E1_2 | | 6.7 | 76.9 | 3.33E+08 | 1.01E+08 | 3324 |
| GA_E2_1 | | 7.5 | 73 | 3.62E+08 | 1.04E+08 | 3927 |
| Precamcrust | Magnabosco et al. (2016)[39] | 4.8 | 95.9 | 8.39E+07 | 5.90E+07 | 6728 |
| Tomsk | Kadnikov et al. (2018)[40] | 18.1 | 98.7 | 1.62E+08 | 1.38E+08 | 26124 |
| AfrMine_BE2011 | Lau et al. (2014)[41] | 4 | 88.1 | 1.39E+08 | 8.53E+07 | 17953 |
| AfrMine_BE2012 | | 4.9 | 84.9 | 2.99E+08 | 1.81E+08 | 9193 |
| AfrMine_DR5 | | 5.2 | 92.6 | 1.24E+08 | 7.67E+07 | 6075 |
| AfrMine_FI88 | | 4.4 | 86.9 | 2.42E+08 | 1.45E+08 | 11589 |
| AfrMine_MM5 | | 2.7 | 95.6 | 4.10E+07 | 2.67E+07 | 5026 |
| AfrMine_TT107 | | 4.7 | 95.9 | 8.34E+07 | 5.86E+07 | 6592 |
| AfrMine_TT109 | | 4.6 | 94.7 | 9.48E+07 | 7.31E+07 | 21066 |
| HURL_140m | Hernsdorf et al. (2017)[42] | 14 | 97.4 | 1.79E+08 | 1.29E+08 | 13736 |
| HURL_250m | | 13.7 | 96 | 2.92E+08 | 1.91E+08 | 15318 |
| IMS | Probst et al. (2014)[12] | 40.3 | 93.4 | 2.41E+08 | 1.13E+08 | 4545 |
| SulCav_AS07-7 | Hamilton et al. (2015)[43] | 79.6 | 95.1 | 1.66E+09 | 4.11E+08 | 6162 |
| SulCav_FS08-3 | | 47.6 | 87.9 | 4.91E+09 | 4.66E+08 | 2219 |
| SulCav_GS09-5 | | 44.8 | 92.8 | 2.68E+09 | 7.70E+08 | 4281 |
| SulCav_PC08-64 | | 37.4 | 92 | 2.47E+09 | 4.69E+08 | 3088 |
| SulCav_PC08-66 | | 36.2 | 87.9 | 3.52E+09 | 6.79E+08 | 2820 |
| SulCav_PC08-3 | | 46.3 | 92.4 | 1.62E+09 | 3.87E+08 | 4522 |
| SulCav_GS10-10 | | 48.3 | 90.9 | 3.70E+09 | 1.02E+09 | 3059 |
| SulCav_FS06-10 | | 76.9 | 96.1 | 1.16E+09 | 3.18E+08 | 3467 |
| Rifle_sed1 | Anantharaman et al. (2016)[44] | 7.2 | 32.4 | 8.40E+08 | 1.94E+08 | 2242 |
| Rifle_sed2 | | 14.6 | 45 | 1.62E+09 | 4.83E+08 | 2305 |
| Rifle_sed3 | | 14.8 | 32.3 | 1.57E+09 | 3.52E+08 | 2078 |
| Rifle_o2low1 | | 41.1 | 82.2 | 3.97E+09 | 2.07E+09 | 4522 |
| Rifle_o2low2 | | 37 | 75.6 | 3.80E+09 | 1.68E+09 | 3594 |
| Rifle_o2low3 | | 4.8 | 44.7 | 6.44E+08 | 1.60E+08 | 2607 |
| Rifle_o2high1 | | 49.5 | 85.7 | 4.26E+09 | 2.08E+09 | 4688 |
| Rifle_o2high2 | | 39.3 | 75.9 | 4.80E+09 | 1.89E+09 | 3564 |
| Rifle_o2high3 | | 37.2 | 71.2 | 4.69E+09 | 1.66E+09 | 3427 |
| CG03_2015 | Probst et al. (2018)[8] | 21.6 | 91.5 | 1.58E+09 | 6.24E+08 | 3987 |

| | | | | | | |
|---|---|---|---|---|---|---|
| CG04_2015 | | 23.1 | 92 | 1.65E+09 | 6.70E+08 | 5131 |
| CG05_2015 | | 20.3 | 91.8 | 1.44E+09 | 5.82E+08 | 4321 |
| CG11_2015 | | 19.2 | 90.6 | 1.48E+09 | 5.83E+08 | 3844 |
| CG12_2015 | | 18.2 | 90.7 | 1.58E+09 | 5.83E+08 | 3948 |
| CG13_2015 | | 17 | 90.7 | 1.37E+09 | 5.04E+08 | 4070 |
| CG21_2015 | | 19.4 | 94.7 | 7.95E+08 | 3.24E+08 | 3743 |
| CG22_2015 | | 23.2 | 93.5 | 1.18E+09 | 5.45E+08 | 4952 |
| CG23_2015 | | 17 | 93.3 | 9.19E+08 | 3.89E+08 | 4222 |

**Table S3 | Recovered genomes, consensus taxonomy, completeness and contamination.** Genome statistics for dereplicated genomes from the Crystal Geyser along with their completeness and contamination are depicted in Probst et al. (2018). Data can be found in TableS3_Genome_information.xlsx

**Table S4 | Origin and accession numbers of Altiarchaeota genomes.**

| Genome | Origin | Accession number |
|---|---|---|
| GA21 | this study | XXXXXXX |
| SulCav | this study | XXXXXXX |
| Alpenaf | this study | XXXXXXX |
| HURL2 | Hernsdorf et al. (2017) | GCA_002841095.1 |
| CG | Probst et al. (2018) | GCA_002789105.1 |
| HURL1 | Hernsdorf et al. (2017) | GCA_002841105.1 |
| SM1 | Probst et al. (2014) | CCXY00000000.1 |
| WOR2 | Bird et al. (2016) | GCA_001723855.1 |
| WORSCG | Bird et al. (2016) | GCA_001723845.1 |
| WOR79 | Bird et al. (2016) | GCA_001723835.1 |
| pMC2 | Rinke et al. (2013) | GCA_000402775.1 |
| IMC4 | Probst et al. (2014) | MCBF00000000.1 |
| HURL3 | Hernsdorf et al. (2017) | GCA_002842715.1 |

**Table S5 | Data on subsurface samples and ecosystems analyzed in this study** containing sampling sites, depth and references. For assembly stats of metagenomes please see Table S2; for statistics of reconstructed genomes please see Table S3.

| Ecosystem | Assembly | Publication of read dataset | Accession number [SRA or MG-RAST] | Depth |
|---|---|---|---|---|
| GA | GA_E1_1 | This study | XXXXXX | 350 m |
| GA | GA_E1_2 | | XXXXXX | |
| GA | GA_E2_1 | | XXXXXX | |
| TT107 | Precamcrust | Magnabosco et al. (2016) | mgm4529964.3 | 3140 m |
| Tomsk | Tomsk | Kadnikov et al. (2018) | SRR7102746 | 2000 m |
| BE | AfrMine_BE2011 | Lau et al. (2014) | mgm4536100.3 | 1339 m |
| BE | AfrMine_BE2012 | | mgm4536472.3 | 1339 m |
| DR5 | AfrMine_DR5 | | mgm4536473.3 | 1046 m |
| FI88 | AfrMine_FI88] | | mgm4536074.3 | 1056 m |
| MM5 | AfrMine_MM5 | | mgm4529965.3 | 1900 m |
| TT107 | AfrMine_TT107 | | mgm4529964.3 | 3048 m |
| TT109 | AfrMine_TT109 | | mgm4536476.3 | 3136 m |
| lHURL | HURL_140m | Hernsdorf et al. (2017) | PRJNA321556 | 140 m |
| dHURL | HURL_250m | | | 250 m |
| IMS | IMS | Probst et al. (2014 | SRR1534154 | 40 m |
| AS | SulCav_AS07-7 | Hamilton et al. (2015) | SRR1559028 | 0 m |
| FS | SulCav_FS08-3 | | SRR1560849 | |
| GS | SulCav_GS09-5 | | SRR1560848 | |
| PC | SulCav_PC08-64 | | SRR1560064 | |
| PC | SulCav_PC08-66 | | SRR1559230 | |
| PC | SulCav_PC08-3 | | SRR1560850 | |
| GS | SulCav_GS10-10 | | SRR1559353 | |
| FS | SulCav_FS06-10 | | SRR1560266 | |
| sed | Rifle_sed1 | Anantharaman et al. (2016) | SRX1085346 | 5 m |
| sed | Rifle_sed2 | | SRX1085347 | |
| sed | Rifle_sed3 | | SRX1085348 | |
| lO2 | Rifle_o2low1 | | SRX1085356 | |
| lO2 | Rifle_o2low2 | | SRX1085358 | |
| lO2 | Rifle_o2low3 | | SRX1085360 | |
| hO2 | Rifle_o2high1 | | SRX1085354 | |
| hO2 | Rifle_o2high2 | | SRX1085350 | |
| hO2 | Rifle_o2high3 | | SRX1085352 | |

| | | | | |
|---|---|---|---|---|
| mCG | CG03_2015 | Probst et al. (2018) | SRS2524942 | 250 m |
| mCG | CG04_2015 | | SRS2524959 | |
| mCG | CG05_2015 | | SRS2524997 | |
| dCG | CG11_2015 | | SRS2525260 | 400 m |
| dCG | CG12_2015 | | SRS2525262 | |
| dCG | CG13_2015 | | SRS2525263 | |
| sCG | CG21_2015 | | SRS2525667 | 150 m |
| sCG | CG22_2015 | | SRS2525670 | |
| sCG | CG23_2015 | | SRS2525668 | |

**Table S6 | Mean and maximum replication indices of bacteria across subsurface ecosystems.** Genomes of samples with biological replicates were dereplicated using dRep and their replication indices were dereplicated by calculating both the mean iRep value across the replicates as well as the maximum iRep value across the replicates. The replication indices are provided in file TableS6_iRep_bacteria.xlsx

**Table S7 | Median iRep values per sample and depth.** Mean iRep for each genome per ecosystem values were used. Samples are ordered by depth. *SulCav*[43]*, IMS*[12]*, AfrMine*[39,41] *and Tomsk*[40] *are publicly available datasets that were leveraged to bin new genomes and published genomes from Rifle*[45]*, HURL*[42] *and CG were used in this study after dereplication using dRep if not already performed on the available genomes.*

| Ecosystem | Sample | Depth | Median iRep |
|---|---|---|---|
| SulCav | AS | 0 | 1.348 |
| SulCav[#] | FS | 0 | 1.395 |
| SulCav* | PC | 0 | 1.473 |
| SulCav[#] | GS | 0 | 1.407 |
| Rifle* | hO2 | 5 | 1.332 |
| IMS | IMS | 40 | 1.336 |
| HURL | sHURL | 140 | 1.411 |
| HURL | dHURL | 250 | 1.344 |
| AfrMine | DR5 | 1050 | 1.276 |
| AfrMine | FI88 | 1060 | 1.290 |
| AfrMine[#] | BE | 1340 | 1.261 |
| AfrMine | MM5 | 1900 | 1.225 |
| Tomsk | Tomsk | 2000 | 1.191 |
| AfrMine[#] | TT107 | 3050 | 1.205 |
| AfrMine | TT109 | 3140 | 1.258 |
| CG* | sCG | 150 | 1.562 |
| CG* | mCG | 300 | 1.527 |
| GA* | GA | 350 | 1.396 |
| CG* | dCG | 400 | 1.540 |
| Rifle* | lO2 | 5 | 1.347 |
| Rifle* | Sed | 5 | 1.588 |

*sample exists as triplicates, grouped for Fig. 3, separate for Fig. 4

[#]sample exists as duplicate, grouped for Fig. 3, separate for Fig. 4

**Table S8 | Pearson Correlations of iRep indices of genomes with specific metabolic capacities across depth.** P-values were adjusted for multiple testing according to the Benjamini-Hochberg method.

**High $CO_2$ ecosystems excluded (referred to in main text)**

| | Adjusted p-values | Correlation-values |
|---|---|---|
| Arsenate reduction/oxidation | 5.68E-08 | -0.42 |
| $C_1$ compounds utilization | 2.59E-10 | -0.45 |
| Carbon fixation | 5.17E-11 | -0.48 |
| Hydrogen metabolism | 3.99E-22 | -0.45 |
| Sulfur oxidation | 2.85E-13 | -0.46 |
| Selenate reduction | 2.15E-05 | -0.48 |
| Halogens breakdown / perchlorate reduction | 2.82E-05 | -0.39 |
| Metals oxidation/reduction | 0.145600971 | -0.37 |
| Nitrogen metabolism | 3.62E-24 | -0.42 |
| Oxygen respiration | 5.84E-26 | -0.43 |
| Sulfur reduction | 0.019817469 | -0.31 |
| Carbon monoxide oxidation | 4.64E-06 | -0.43 |
| Urea utilization | 0.19515825 | -0.27 |
| Methane oxidation | 0.141004585 | -0.98 |

**High $CO_2$ ecosystems included**

| | Adjusted p-values | Correlation-values |
|---|---|---|
| Arsenate reduction/oxidation | 2.27E-08 | -0.35 |
| $C_1$ compounds utilization | 3.89E-12 | -0.41 |
| Carbon fixation | 9.25E-10 | -0.39 |
| Hydrogen metabolism | 3.81E-29 | -0.45 |
| Sulfur oxidation | 3.05E-13 | -0.46 |
| Selenate reduction | 7.24E-06 | -0.43 |
| Halogens breakdown / perchlorate reduction | 3.00E-06 | -0.36 |
| Metals oxidation/reduction | 0.055037708 | -0.33 |
| Nitrogen metabolism | 4.62E-26 | -0.38 |
| Oxygen respiration | 9.04E-29 | -0.39 |
| Sulfur reduction | 0.021233003 | -0.31 |
| Carbon monoxide oxidation | 4.45E-07 | -0.41 |
| Urea utilization | 0.052759627 | -0.33 |
| Methane oxidation | 0.129494006 | -0.98 |

**Table S9 | Two-group significance (high vs non-high $CO_2$ ecosystems) testing results of metabolic potentials in assemblies.** The read-normalized abundances of metabolic pathways of high $CO_2$ ecosystems were compared using Welch's t-test and the Kruskal-Wallis test. P-values were adjusted using the Benjamini-Hochberg method.

| **Metabolism** | **t-test 2-independent group P-values** | **Kruskal-Wallis group comparison** |
|---|---|---|
| Hydrogen metabolism | 0.86 | 0.62 |
| Oxygen respiration | 0.28 | 0.23 |
| Sulfur metabolism | 0.86 | 0.97 |
| rTCA | 0.86 | 0.23 |
| Carbon monoxide oxidation | 0.86 | 0.11 |
| Wood Lijungdahl pathway | 0.86 | 0.38 |
| Formaldehyde oxidation | 0.27 | 0.23 |
| Formate oxidation | 0.86 | 0.85 |
| Methylamine to formaldehyde | 0.86 | 0.85 |
| Methanol oxidation | 0.97 | 0.39 |
| Nitrate reduction | 0.86 | 0.38 |
| $N_2$ fixation | 0.48 | 0.67 |
| Nitric oxide reduction | 0.86 | 0.67 |
| Nitrite reduction | 0.02 | 0.0006 |
| Nitrous oxide reduction | 0.28 | 0.62 |
| Methane oxidation (PMO) | 0.27 | 0.06 |
| CBB pathway | 0.27 | 0.85 |
| Nitrite oxidation | 0.44 | 0.05 |

**Additional supplementary files**

**File 1 | Tree file of all bacterial genomes using 16 ribosomal proteins**: FileS1_Bornemann et_al_mantle_degassing.tree

**File 2 | Altiarchaeles 16S rRNA gene-based phylogeny**. One representative of each DPANN phylum was used as the outgroup: FileS2_Bornemann et_al_mantle_degassing.newick

**File 3 | Additional geochemical measurements of various laboratories from 1904 to 2004.** Measurements were done according to the german TrinwV-GW (drinking water guidelines). The additional geochemical measurements are provided in file FileS3_Bornemann et_al_mantle_degassing_bioRxiv.xlsx

### References


1. Whitman, W. B., Coleman, D. C. & Wiebe, W. J. Prokaryotes: The unseen majority. *Proc. Natl. Acad. Sci.* **95**, 6578–6583 (1998).
2. Hug, L. A. *et al.* A new view of the tree of life. *Nat. Microbiol.* **1**, 16048 (2016).
3. Dick, G. J. *et al.* Community-wide analysis of microbial genome sequence signatures. *Genome Biol.* **10**, R85 (2009).
4. Brown, C. T., Olm, M. R., Thomas, B. C. & Banfield, J. F. Measurement of bacterial replication rates in microbial communities. *Nat. Biotechnol.* **34**, 1256 (2016).
5. Wu, Y.-W., Simmons, B. A. & Singer, S. W. MaxBin 2.0: an automated binning algorithm to recover genomes from multiple metagenomic datasets. *Bioinformatics* **32**, 605–607 (2016).
6. Alneberg, J. *et al.* Binning metagenomic contigs by coverage and composition. *Nat. Methods* **11**, 1144–1146 (2014).
7. Sieber, C. M. K. *et al.* Recovery of genomes from metagenomes via a dereplication, aggregation and scoring strategy. *Nat. Microbiol.* **3**, 836–843 (2018).
8. Probst, A. J. *et al.* Differential depth distribution of microbial function and putative symbionts through sediment-hosted aquifers in the deep terrestrial subsurface. *Nat. Microbiol.* **3**, 328–336 (2018).
9. Olm, M. R., Brown, C. T., Brooks, B. & Banfield, J. F. dRep: a tool for fast and accurate genomic comparisons that enables improved genome recovery from metagenomes through de-replication. *ISME J.* **11**, 2864 (2017).
10. Cline, J. D. Spectrophotometric Determination of Hydrogen Sulfide in Natural Waters1. *Limnol. Oceanogr.* **14**, 454–458 (1969).
11. Moissl, C., Rachel, R., Briegel, A., Engelhardt, H. & Huber, R. The unique structure of archaeal 'hami', highly complex cell appendages with nano-grappling hooks: Unique structure of archaeal 'hami'. *Mol. Microbiol.* **56**, 361–370 (2005).
12. Probst, A. J. *et al.* Biology of a widespread uncultivated archaeon that contributes to carbon fixation in the subsurface. *Nat. Commun.* **5**, 5497 (2014).
13. Bird, J. T., Baker, B. J., Probst, A. J., Podar, M. & Lloyd, K. G. Culture Independent Genomic Comparisons Reveal Environmental Adaptations for Altiarchaeales. *Front. Microbiol.* **7**, (2016).
14. Papke, R. T., Ramsing, N. B., Bateson, M. M. & Ward, D. M. Geographical isolation in hot spring cyanobacteria. *Environ. Microbiol.* **5**, 650–659 (2003).
15. Whitaker, R. J., Grogan, D. W. & Taylor, J. W. Geographic barriers isolate endemic populations of hyperthermophilic archaea. *Science* **301**, 976–978 (2003).
16. Liu, L. *et al.* High correlation between genotypes and phenotypes of environmental bacteria Comamonas testosteroni strains. *BMC Genomics* **16**, (2015).
17. Wright, S. Isolation by Distance. *Genetics* **28**, 114–138 (1943).
18. Wright, S. Isolation by Distance Under Diverse Systems of Mating. *Genetics* **31**, 39–59 (1946).
19. Malecot, G. Mathematics of heredity. *Math. Hered.* (1948).
20. Kimura, M. & Weiss, G. H. The Stepping Stone Model of Population Structure and the Decrease of Genetic Correlation with Distance. *Genetics* **49**, 561–576 (1964).
21. Maruyama, M. On automorphism groups of ruled surfaces. *J. Math. Kyoto Univ.* **11**, 89–112 (1971).

22. Martiny, J. B. H. *et al.* Microbial biogeography: putting microorganisms on the map. *Nat. Rev. Microbiol.* **4**, 102–112 (2006).
23. Hanson, C. A., Fuhrman, J. A., Horner-Devine, M. C. & Martiny, J. B. H. Beyond biogeographic patterns: processes shaping the microbial landscape. *Nat. Rev. Microbiol.* **10**, 497–506 (2012).
24. Gaisin, V. A. *et al.* Biogeography of thermophilic phototrophic bacteria belonging to Roseiflexus genus. *FEMS Microbiol. Ecol.* **92**, (2016).
25. Varliero, G., Bienhold, C., Schmid, F., Boetius, A. & Molari, M. Microbial Diversity and Connectivity in Deep-Sea Sediments of the South Atlantic Polar Front. *Front. Microbiol.* **10**, (2019).
26. Cocks, L. R. M. & Torsvik, T. H. Baltica from the late Precambrian to mid-Palaeozoic times: The gain and loss of a terrane's identity. *Earth-Sci. Rev.* **72**, 39–66 (2005).
27. Torsvik, T. H. *et al.* Phanerozoic polar wander, palaeogeography and dynamics. *Earth-Sci. Rev.* **114**, 325–368 (2012).
28. Maruyama, S., Isozaki, Y., Kimura, G. & Terabayashi, M. Paleogeographic maps of the Japanese Islands: Plate tectonic synthesis from 750 Ma to the present. *Isl. Arc* **6**, 121–142 (1997).
29. Lovley, D. R. & Chapelle, F. H. Deep subsurface microbial processes. *Rev. Geophys.* **33**, 365–381 (1995).
30. Magnabosco, C. *et al.* The biomass and biodiversity of the continental subsurface. *Nat. Geosci.* **11**, 707–717 (2018).
31. Pebesma, E. & Bivand, R. Classes and Methods for Spatial Data in R. *R News* **5**, (2005).
32. Jain, C., Rodriguez-R, L. M., Phillippy, A. M., Konstantinidis, K. T. & Aluru, S. High throughput ANI analysis of 90K prokaryotic genomes reveals clear species boundaries. *Nat. Commun.* **9**, (2018).
33. Hoang, D. T., Chernomor, O., von Haeseler, A., Minh, B. Q. & Vinh, L. S. UFBoot2: Improving the Ultrafast Bootstrap Approximation. *Mol. Biol. Evol.* **35**, 518–522 (2018).
34. Guindon, S. *et al.* New algorithms and methods to estimate maximum-likelihood phylogenies: assessing the performance of PhyML 3.0. *Syst. Biol.* **59**, 307–321 (2010).
35. Anisimova, M., Gil, M., Dufayard, J.-F., Dessimoz, C. & Gascuel, O. Survey of Branch Support Methods Demonstrates Accuracy, Power, and Robustness of Fast Likelihood-based Approximation Schemes. *Syst. Biol.* **60**, 685–699 (2011).
36. Oksanen, J. *et al.* vegan: Community Ecology Package. (2012).
37. Shannon, P. *et al.* Cytoscape: a software environment for integrated models of biomolecular interaction networks. *Genome Res.* **13**, 2498–2504 (2003).
38. Bräuer, K., Kämpf, H., Niedermann, S. & Strauch, G. Indications for the existence of different magmatic reservoirs beneath the Eifel area (Germany): A multi-isotope (C, N, He, Ne, Ar) approach. *Chem. Geol.* **356**, 193–208 (2013).
39. Magnabosco, C. *et al.* A metagenomic window into carbon metabolism at 3 km depth in Precambrian continental crust. *ISME J.* **10**, 730 (2016).
40. Kadnikov, V. V. *et al.* A metagenomic window into the 2-km-deep terrestrial subsurface aquifer revealed multiple pathways of organic matter decomposition. *FEMS Microbiol. Ecol.* **94**, (2018).
41. Lau, M. C. Y. *et al.* Phylogeny and phylogeography of functional genes shared among seven terrestrial subsurface metagenomes reveal N-cycling and microbial evolutionary relationships. *Front. Microbiol.* **5**, (2014).

42. Hernsdorf, A. W. *et al.* Potential for microbial $H_2$ and metal transformations associated with novel bacteria and archaea in deep terrestrial subsurface sediments. *ISME J.* **11**, 1915–1929 (2017).
43. Hamilton, T. L., Jones, D. S., Schaperdoth, I. & Macalady, J. L. Metagenomic insights into S(0) precipitation in a terrestrial subsurface lithoautotrophic ecosystem. *Front. Microbiol.* **5**, (2015).
44. Anantharaman, K. *et al.* Thousands of microbial genomes shed light on interconnected biogeochemical processes in an aquifer system. *Nat. Commun.* **7**, (2016).
